# Mechanistic Insights into TMPyP4 Recognition of the HIV-1 LTR-III G-Quadruplex in Dilute and Protein Condensate Environments Reveal Hidden Dual Binding Modes

**DOI:** 10.64898/2026.05.16.724744

**Authors:** Susmita Pradhan, Shailesh Mani Tripathi, Shubhangi Sharma, Arun Pratap Singh, Sandeep Sundriyal, Satyajit Patra

## Abstract

G-quadruplex (GQ) structures within the HIV-1 long terminal repeat (LTR) regulate viral transcription and represent promising antiviral targets; however, detailed mechanistic understanding of their ligand recognition at the molecular level remains limited and has largely been investigated under dilute conditions despite the crowded and compartmentalized nature of intracellular environment. Here, we investigate the interaction of the cationic porphyrin TMPyP4 with the HIV-1 LTR-III GQ under dilute conditions and inside protein-rich phase-separated condensates that mimic intracellular biocondensates. Steady-state and time-resolved fluorescence measurements reveal a dual binding behavior that is not discernible from absorption spectroscopy. A high-affinity guanine-rich binding mode leads to efficient fluorescence quenching through electron transfer from ground-state guanine to excited TMPyP4, whereas a weaker non-guanine binding mode gives rise to enhanced and long-lived emission. Nucleotide-specific control experiments validate the origin of these distinct binding environments. Molecular docking and molecular dynamics simulations further support preferential binding of TMPyP4 at the terminal G-quartet together with a secondary binding mode near the quadruplex–duplex junction. Importantly, both TMPyP4 and LTR-III GQ preferentially partition into the condensates, where the hybrid GQ structure, dual binding behavior, and associated excited-state signatures remain preserved despite the crowded and viscous environment. Although a slight reduction in binding affinity is observed inside the condensates, the overall binding mechanism remains largely preserved due to compensatory effects arising from the condensate microenvironment. Overall, this work demonstrates that ligand recognition of viral GQ remains preserved within protein condensates and establishes fluorescence spectroscopy as a sensitive approach for resolving hidden binding heterogeneity in GQ–ligand interactions.

## Introduction

Human immunodeficiency virus type-1 (HIV-1) remains a major global health challenge, and despite the success of antiretroviral therapy, the absence of a definitive cure necessitates the identification of new molecular targets for antiviral intervention. The long terminal repeat (LTR) region of the HIV-1 genome plays a central role in regulating viral transcription and contains conserved guanine-rich sequences capable of forming G-quadruplex (GQ) structures under physiological conditions.^1–4^ These GQ structures act as regulatory elements, where their stabilization suppresses transcription by inhibiting transcription factor binding, while their disruption enhances viral expression.^3,5,6^ Consequently, the LTR GQ has emerged as a promising target for therapeutic intervention.^4,7–12^

Among the GQ motifs present in the LTR, the LTR-III sequence adopts a unique quadruplex–duplex hybrid structure, as revealed by NMR studies, consisting of a (3 + 1) GQ core coupled to a duplex stem through a junction region.^13^ This structural organization introduces distinct binding environments, particularly at the quadruplex–duplex junction, which provides opportunities for selective ligand recognition.^4^ A wide range of small molecules have been designed to target this structure, and their equilibrium binding affinities have been extensively characterized under dilute conditions.^7,10,14^ However, detailed mechanistic understanding of ligand recognition of the HIV-1 LTR-III GQ at the molecular level—particularly the existence of multiple binding modes and their origin within different regions of the GQ scaffold—remains largely understudied. Most studies of GQ–ligand interactions have been carried out under dilute solution conditions, which do not capture the crowded and compartmentalized intracellular environment. In addition to membrane-bound organelles, biomolecules are also organized into dynamic membraneless compartments formed via liquid–liquid phase separation (LLPS), such as nucleoli, stress granules, and Cajal bodies. These condensates arise from multivalent interactions between biomolecules and generate microenvironments that differ significantly from dilute solutions in terms of viscosity, hydration, dielectric properties, and molecular mobility.^15–18^ Such changes are expected to influence both biomolecular structure and ligand binding interactions.^15,19–28^ While significant progress has been made in understanding the thermodynamic and physicochemical properties of these condensates,^29,30^ how such environments influence both G-quadruplex structure and its recognition by small-molecule ligands at a molecular level remains largely unexplored.^21^

TMPyP4 is a well-characterized GQ-binding ligand whose fluorescence is highly sensitive to the local environment and binding mode, making it suitable for probing binding heterogeneity.^31–35^ To mimic condensate-like environments, we employ a BSA–PEG aqueous two-phase system (ATPS) that forms protein-rich droplets with well-defined physicochemical properties, providing a suitable platform to examine the effects of crowding, reduced water activity, and increased viscosity on biomolecular interactions.^36–38^

In this work, we investigate the interaction of TMPyP4 with the HIV-1 LTR-III GQ in both buffer and protein-rich condensates formed in a bovine serum albumin (BSA)–polyethylene glycol (PEG) aqueous two-phase system (ATPS). By combining steady-state and time-resolved fluorescence spectroscopy, we uncover a dual binding behavior that is not evident from absorption measurements, arising from distinct interactions at guanine-rich and non-guanine regions of the GQ. Nucleotide-specific control experiments, together with molecular docking and molecular dynamics simulations, are used to assign these binding modes. Furthermore, we demonstrate that both the structure of the GQ and its mode of ligand recognition are preserved within condensate-like environments, despite significant changes in physicochemical conditions. This work provides mechanistic insight into GQ–ligand interactions at the molecular level and establishes that the key features of ligand recognition remain preserved under biologically relevant crowded conditions.

## 2. Experimental Sections

### 2.1. Materials and Sample Preparation

Three types DNA oligonucleotides were used in this study. The sequences of the DNA oligonucleotides are the following:

**LTR-III GQ**: 5’-GG GA GGC GTG GCC T GGG C GGG ACT GGG G-3’

**Truncated LTR-III GQ**: 5’-GG GA T GGG C GGG ACT GGG G-3’

**Duplex Loop**: 5’-GGC GTG GCC-3’

In the case of truncated LTR-III GQ, the duplex hairpin forming sequence is removed. For duplex loop sequence only the duplex hairpin sequences were chosen. All the DNA oligonucleotides were purchased from Integrated DNA Technologies (IDT). Prior to use all the nucleotides were rapidly heated at 90^0^C for 3 minutes followed by gradually cooled to room temperature over 2-3 hours in the experimental buffer containing 30 mM Tris-HCl, 100 mM NaCl at pH 7.4 to ensure proper folding of the DNA oligonucleotides. Tris(hydroxymethyl) aminomethane (Tris) ≥ 99.9%, hydrochloric acid (HCl, 37%, ACS reagent), sodium chloride (NaCl, ACS reagent, ≥ 99.0%) for the preparation of buffer were procured from Sigma Aldrich (Merck). Polyethylene glycol (PEG) average M_n_ 4.6 kDa (373001), bovine serum albumin (BSA, A3059), *meso*-5,10,15,20-Tetrakis-(N-methyl-4-pyridyl) porphine tetratosylate (TMPyP4) were also purchased from Sigma Aldrich (Merck). Alexa 647 NHS ester (A20006) was obtained from ThermoFisher Scientific. All the experiments were performed at 25^0^C. MilliQ water is used in all the measurements.

### 2.2. Methods

#### 2.2.1. Cloud Point Titration

The binodal curve for the BSA-PEG aqueous two-phase system was determined from the turbidometric measurements. To determine the turbidity of the aqueous mixture of BSA and PEG cloud point titration was carried out. For this purpose, the absorption of the BSA-PEG aqueous solution was measured at 350 nm in a Nanodrop UV-Visible spectrophotometer. The total concentration of BSA and PEG was remained constant at 240 mg/ml and their relative ratio, [BSA]/[PEG] was varied from 0.04 to 1.8.

#### 2.2.2. UV-Visible Absorption Measurements

A V-650 (JASCO) spectrophotometer was used for all the absorption measurements. A microfluorimeter cell of path length 1 cm and volume 700 μL (18F-Q-10, Starna) was used to obtain the absorption spectrum of TMPyP4 as a function of LTR-III GQ concentrations. In the spectrophotometric titration the TMPyP4 concentration was fixed at 2 μM and the GQ concentrations were varying from 0 μM to 30 μM. In the case of Job’s plot measurement, the total concentration of LTR-III GQ and TMPyP4 was fixed at 50 μM and mole fractions of TMPyP4 were varying from 0.1 to 1.0.

#### 2.2.3. Steady State Fluorescence Measurement

Steady-state fluorescence measurements were performed using Fluorolog (Horiba) and Fluoromax-4 (Horiba) spectrofluorimeters. Fluorescence spectra of TMPyP4 were recorded in a quartz microfluorimeter cuvette (18F-Q-10, Starna). During fluorescence titration experiments, the concentration of TMPyP4 was fixed at 2 μM, while the concentration of LTR-III G-quadruplex (GQ) was varied from 0 to 80 μM. TMPyP4 was excited at 434 nm, and the excitation and emission slit widths were set to 2 nm and 5 nm, respectively. Fluorescence spectra were recorded as a function of GQ concentration, and integrated fluorescence intensities were obtained by integrating the area under the emission spectrum at each titration point. These integrated intensities were plotted as a function of GQ concentration to generate binding curves.

The binding curves were analysed using an independent and equivalent binding sites model (**Section 1.1, Supporting Information**). For the low-GQ-concentration region, the fluorescence quenching data were fitted using

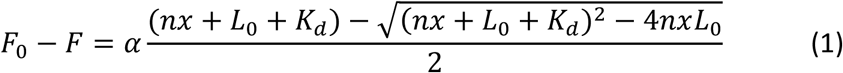

where *F* and *F*_0_ are the fluorescence intensities of TMPyP4 in the presence and absence of GQ, respectively. Here, *n* is the number of binding sites per GQ, *x* is the total GQ concentration, *L*_0_ is the total TMPyP4 concentration, *K*_*d*_ is the microscopic dissociation constant for a single binding site, and *α* is the proportionality constant relating the fluorescence response to the concentration of bound TMPyP4. For the high-GQ-concentration region, the fluorescence enhancement data were fitted using

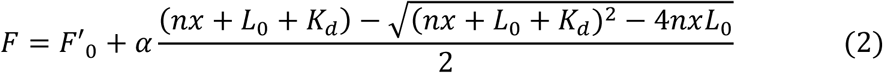

where *F*represents the fluorescence intensity at different GQ concentrations in the fluorescence-enhancement regime, and 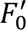 corresponds to the minimum fluorescence intensity obtained during the titration, i.e., the fluorescence intensity immediately before the onset of fluorescence enhancement. All nonlinear least-squares fitting was performed using OriginPro 2023 employing the Levenberg–Marquardt algorithm.

#### 2.2.4. Time Resolved Fluorescence Measurement

Time-resolved fluorescence measurements were performed using time-correlated single-photon counting (TCSPC) spectrometers (DeltaFlex-01, Horiba). Lifetime measurements in buffer were carried out using the DeltaFlex TCSPC facility at SATHI, IIT Delhi (Sonipat campus), while measurements in protein droplets were conducted using a DeltaFlex-01 TCSPC system at the Central Analytical Laboratory (CAL), BITS Pilani. For measurements in buffer, TMPyP4 was excited using a 440 nm pulsed laser diode operating at a repetition rate of 1 MHz, and fluorescence decays were collected at 660 nm as a function of GQ concentration. For measurements in protein droplets, excitation was performed using a 455 nm pulsed light-emitting diode (DeltaDiode-455) operating at 1 MHz, and emission decays were recorded at 660 nm. Photon detection was achieved using a picosecond photon detection module (PPD-850) equipped with a fast, cooled photomultiplier tube. The instrument response function (IRF) was determined using a dilute aqueous suspension of Ludox as a scattering standard. All decay curves were analyzed using EzTime software employing iterative reconvolution.

The fluorescence decay profiles were analyzed using multiexponential functions of the form:

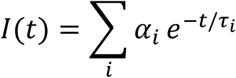

where *τ*_*i*_ represents the lifetime of the *i*th component and *α*_*i*_ is the corresponding normalized pre-exponential amplitude, with ∑_*i*_ *α*_*i*_ = 1. For TMPyP4 in buffer in the absence of GQ, the decay curves were adequately described by a biexponential function. In all other cases, a triexponential model was required to achieve optimal fits, as determined from reduced chi-square (χ²) values close to unity and the random distribution of residuals. The relative intensity fraction (*f*_*i*_) of each lifetime component was calculated as: 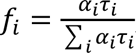, which represents the fractional contribution of each component to the total time-integrated fluorescence intensity. The amplitude-weighted and intensity-weighted average lifetimes were calculated using:

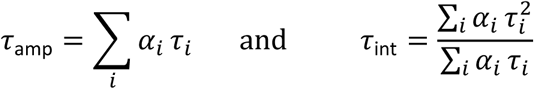

#### 2.2.5. Circular Dichroism (CD) Studies

Circular dichroism (CD) measurements were performed to record and compare CD spectra of HIV-1 LTR-III G-quadruplex (GQ) in buffer and in BSA droplets, both in the absence and presence of TMPyP4. CD spectra were recorded at 25^0^C using a BioLogic MOS-500 spectrometer at the Central Research Facilities (CRF), IIT Delhi (Sonipat campus). Spectra were collected over the wavelength range 190–400 nm using an 800 μL quartz cuvette with a path length of 0.2 cm. In all CD measurements, the concentration of LTR-III GQ was fixed at 10 μM. For measurements in the presence of TMPyP4, TMPyP4 was added to a final concentration of 5 μM to the solution containing 10 μM GQ. Protein droplets were prepared by forming an aqueous two-phase system consisting of 170 mg mL⁻¹ polyethylene glycol (PEG) and 70 mg mL⁻¹ bovine serum albumin (BSA).

#### 2.2.6. Phase Contrast, Differential Interference Contrast (DIC), and Fluorescence Confocal Imaging of the BSA droplets

A Zeiss apotome microscope was used for the phase contrast imaging of the BSA droplets. Fluorescence and DIC imaging of the BSA droplets were conducted on a confocal laser scanning fluorescence microscope (Zeiss LSM 880). To image the alexa 647 labelled BSA droplets, the samples were excited at 633 nm. ATPS samples with LTR-III GQ and TMPyP4 were excited at 440 nm. To focus the excitation light on the sample a 10x plan apochromat objective lens, NA 0.45 (Zeiss) was used.

#### 2.2.7. Fluorescent labelling of BSA with Alexa Fluor 647

Fluorescent labelling of bovine serum albumin (BSA) was performed using Alexa Fluor 647 NHS ester via standard amine-reactive chemistry.^39,40^ Briefly, BSA (∼30 µM) in 0.2 M NaHCO₃ buffer (pH 9.2) was reacted with Alexa Fluor 647 (8 mM stock in DMSO) by adding 20 µL of the dye solution to 1.98 mL of BSA solution, giving a final dye concentration of ∼80 µM (≈3-fold molar excess over BSA). The reaction was carried out in the dark for 2 h with gentle stirring. The mixture was then purified using a Sephadex G-25 size-exclusion column equilibrated with phosphate buffer (pH 7.4) to remove unreacted dye. Fractions containing the BSA–fluorophore conjugate were identified by their absorption profile, and the degree of labelling was determined from absorbance at 651 nm and 280 nm using the respective extinction coefficients.

#### 2.2.8. Molecular Docking and Molecular Dynamics Simulation studies

Molecular docking and molecular dynamics (MD) simulations were performed to investigate possible binding modes of TMPyP4 with the HIV-1 LTR-III G-quadruplex (GQ). The G-quadruplex structure of HIV LTR-III (**PDB ID: 6H1K**) was obtained from the RCSB Protein Data Bank.^13^ As the deposited structure lacked coordinated K^+^ ions within the central channel, two K^+^ ions were introduced between adjacent G-quartets to generate an ion-stabilized GQ structure. The resulting structure was subjected to an initial 200 ns MD simulation, and the representative structure from the most populated conformational cluster was used for subsequent docking studies. Prior to docking, the representative GQ structure was prepared using the Protein Preparation Workflow in Schrödinger suite (v2025-4),^41^ including hydrogen addition, optimization of hydrogen-bonding networks, and energy minimization using the OPLS4 force field.^42^ The geometry of the TMPyP4 ligand was optimized at the HF/6-31G* level of theory using Gaussian 09 to obtain the energy minimized conformation for docking. Electrostatic potential (ESP) charges were subsequently calculated at the same level of theory. Molecular docking of TMPyP4 was carried out employing Glide program.^43,44^ Multiple receptor grids were generated covering different structural regions of the LTR-III GQ, including terminal G-quartets, the quadruplex–duplex junction, and duplex loop regions. Docked poses were evaluated based on docking score, MM-GBSA binding free energy, and π–π stacking interactions with guanine-rich regions, and representative binding modes were selected for further MD simulations.

To evaluate the dynamic stability of the modelled binding modes, explicit solvent MD simulations were performed on three distinct systems: the apo ion-stabilized LTR-III GQ (simulated for 200 ns) and two DNA-TMPyP4 docked complexes (simulated for 500 ns each). Restrained electrostatic potential (RESP) charge fitting of TMPyP4 was performed using the antechamber module of AmberTools 18.^45^ The DNA was parameterized using the AMBER DNA OL15 force field.^46,47^ To accurately capture the critical ion-nucleic acid interaction, all K^+^ ions, both channel-coordinated and bulk-solvated, were parametrized using the K^+^ parameters developed by the Cheatham group.^48^ System preparation in Leap involved solvation in a TIP3P water model with a truncated octahedral box and a 10 Å buffer between the solute and box edges. Following neutralization with K^+^ counterions, 100 mM KCl was added to the system to mimic the physiological ionic strength.^49^

Each complex underwent 10,000 steps of restrained minimization using the steepest descent method with a restraint force constant of 2.0 kcal mol^-1^ Å^-2^. This was followed by 300 ps of heating, where the temperature was ramped from 0 to 300 K over 100 ps, and then 200 ps of simulation at a target temperature of 300 K using a Langevin thermostat. During heating, restraints (2 kcal mol^-1^ Å^-2^) were applied to the DNA and two K^+^ ions, followed by 200 ps of constant volume, constant temperature (NVT) ensemble simulation. Next, the system was equilibrated for 200 ps under constant pressure (NPT) using a Berendsen barostat, with restraints gradually reduced from 2.0 to 0.0 kcal mol^-1^ Å^-2^ over 1800 ps. Finally, a production run was conducted at constant temperature and pressure with periodic boundary conditions (PBC) and without restraints, using the GPU-accelerated PMEMD engine in AMBER 18.^50^

The SHAKE algorithm was applied to constrain bond lengths involving hydrogen atoms, with a 2.0 fs timestep.^51^ Long-range electrostatics under PBC were computed using the particle-mesh Ewald (PME) method, and a 10 Å cutoff was applied for short-range nonbonded interactions, with van der Waals interactions treated via a uniform density approximation.^52^ MD trajectory analysis was performed using the CPPTRAJ module of AmberTools18. For visualization purposes, VMD, ChimeraX, Discovery Studio Visualizer and PyMOL software were employed to examine and interpret the structures of interest.^53^ Binding free energies of the TMPyP4–GQ complexes were estimated using the MM-PBSA approach implemented in MMPBSA.py according to: Δ*G*_*bind*_ = *G*_*complex*_ − (*G*_*GQ*_ + *G*_*ligand*_), where *G*_*complex*_, *G*_*GQ*_, and *G*_*ligand*_ represent the free energies of the complex, isolated GQ, and ligand, respectively. Binding free energies were calculated from snapshots extracted from the production trajectories.^54^

## 3. Results and Discussions

### 3.1. Mechanistic insights into the binding interaction in buffer

#### 3.1.1. Steady state absorption and fluorescence measurements

To obtain mechanistic insight into the interaction between TMPyP4 and HIV-1 LTR-III GQ in buffer, binding titrations were performed using steady-state UV–Vis absorption and fluorescence spectroscopy, with TMPyP4 fixed at 2 μM (**Figure 1A,B**). Upon addition of GQ, the Soret band at 422 nm exhibits pronounced hypochromicity along with a bathochromic shift to 440 nm, with a well-defined isosbestic point at 434 nm (**Figure 1A**), indicating clean interconversion between free and bound species. The large decrease in absorbance together with the ∼19 nm red shift is characteristic of strong π–π interactions and is consistent with end-stacking of TMPyP4 onto the terminal G-quartet.^31,34,55^ The absorbance changes reach saturation at ∼5 μM GQ, indicating that most TMPyP4 molecules are bound under these conditions, with negligible further perturbation of the ground-state electronic structure at higher GQ concentrations. The binding stoichiometry was examined using a Job plot at a fixed total concentration of 50 μM (**Figure 1A**, bottom), where the maximum appears at x_TMPyP4_ ≈ 0.6–0.7, consistent with a predominant 2:1 TMPyP4:GQ complex. It is important to note that absorption measurements here primarily reflect the dominant interaction and do not distinguish between different binding modes.

**Figure 1.**
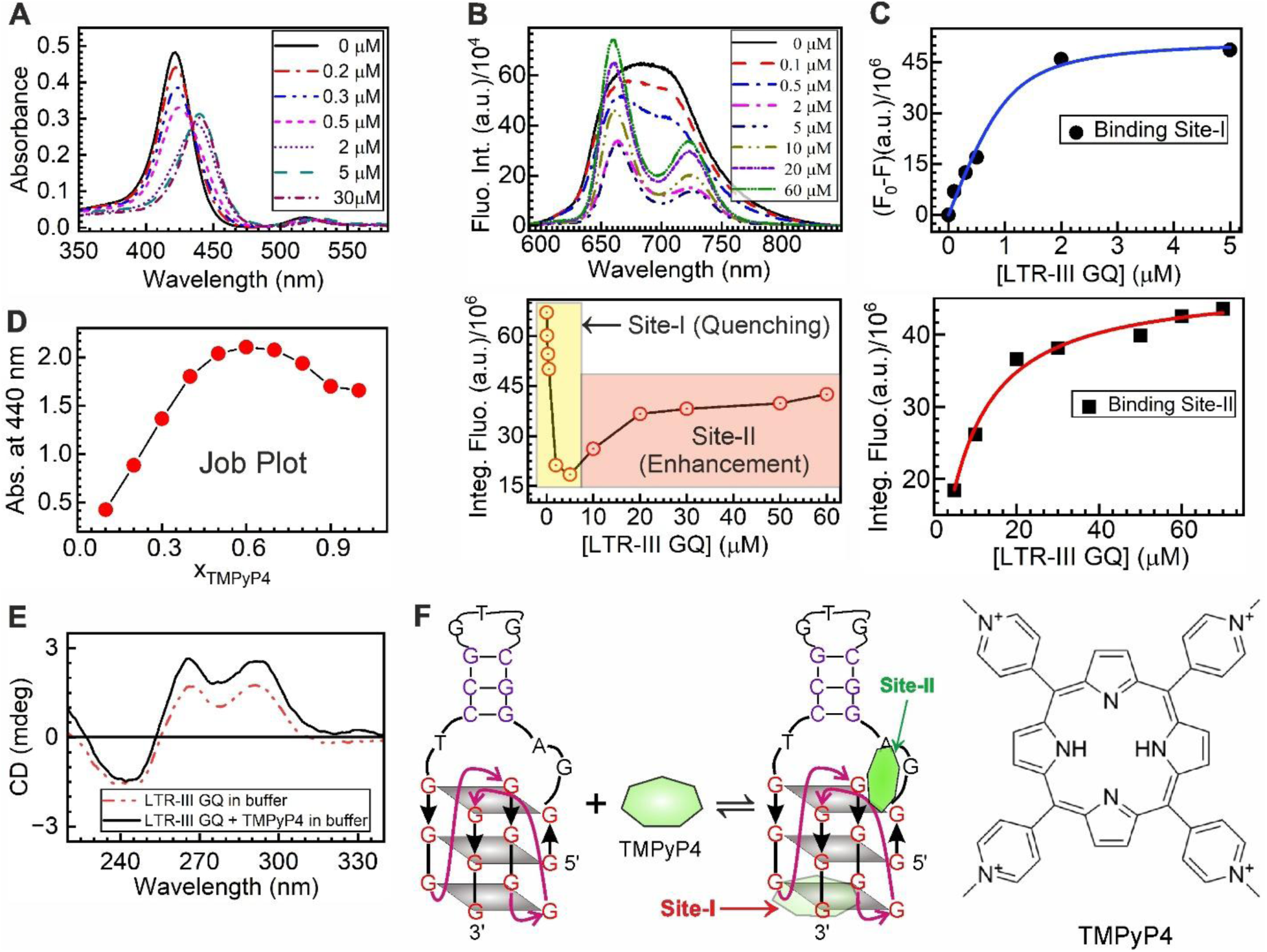
Binding titration of TMPyP4 with the HIV-1 LTR-III GQ in buffer. (A) Absorption spectra of TMPyP4 upon titration with increasing concentrations of LTR-III GQ. The Soret band at 421 nm decreases while a new band at 440 nm grows, with an isosbestic point at ∼434 nm, indicating complex formation. (B) Fluorescence spectra of TMPyP4 as a function of GQ concentration. The fluorescence intensity decreases up to ∼5 μM GQ, followed by an increase at higher concentrations (**Top Panel**). The broad emission profile becomes progressively structured, with two peaks clearly resolved from ∼2 μM GQ onward. Integrated fluorescence intensity as a function of GQ concentration (**Bottom panel**), showing two distinct binding regimes: an initial quenching phase (Site-I) followed by fluorescence enhancement (Site-II) at higher concentrations. (C) Binding isotherm obtained from the quenching regime (0–5 μM GQ). ΔF = (F₀ − F), where F₀ is the integrated fluorescence intensity of free TMPyP4 and F is that in the presence of GQ. The curve saturates near ∼2 μM GQ and was fitted using Equation 1 to determine the dissociation constant (K_d_). (E) Binding isotherm from the fluorescence enhancement regime (>5 μM GQ), plotted as integrated fluorescence intensity versus GQ concentration. The data were fitted with **Equation 2** to extract the corresponding K_d_ value.

Further insight into the binding interaction was obtained from steady-state fluorescence measurements under identical titration conditions (**Figure 1B**). A distinct bimodal fluorescence response is observed, where the fluorescence intensity decreases sharply at low GQ concentrations (0–5 μM) and then increases at higher concentrations, reaching saturation at ∼40 μM (**Figure 1B**, bottom). This behavior indicates the presence of two binding environments with markedly different excited-state deactivation pathways.^34^ In aqueous buffer, TMPyP4 exhibits a broad and featureless emission band, which resolves into two distinct peaks centered at ∼660 nm and ∼723 nm upon binding. This behavior arises from coupling between the first excited singlet (S₁) state and a nearby charge-transfer (CT) state involving intramolecular electron transfer from the porphyrin core to the pyridinium substituents.^56^ In aqueous solution, this coupling is enhanced by the high polarity of the solvent and conformational flexibility of TMPyP4, whereas upon binding to GQ, reduced polarity and restricted rotational freedom weaken the S₁–CT coupling, leading to the resolution of the emission into two distinct bands.^31,35,56^ Notably, in the fluorescence enhancement regime, the relative intensity of the ∼660 nm band increases, indicating a growing contribution from a binding environment where quenching is suppressed.

The integrated fluorescence intensity at each GQ concentration was used to construct binding curves for the two regimes (**Figure 1C**), where the quenching (Site-I) and enhancement (Site-II) regions were analyzed separately using the same binding model, as described in the Experimental section. The resulting dissociation constants (K_d_) are 408.7 ± 50.0 nM for Site-I and 6.7 ± 0.7 μM for Site-II, indicating that Site-I binds approximately one order of magnitude more strongly than Site-II, consistent with previous reports.^55^ To probe the chemical origin of these binding environments, control experiments with individual nucleotides (AMP, GMP, TMP, and CMP) were performed (**Figure S1**), where fluorescence quenching is observed only in the presence of GMP, whereas other nucleobases induce fluorescence enhancement. This behavior arises from electron transfer from the ground state of guanine to the excited state of TMPyP4, which is thermodynamically favorable because the excited-state reduction potential of TMPyP4 (>1.54 V) is higher than that of guanine (∼1.47 V),^33,57^ while the corresponding reduction potentials of AMP, TMP, and CMP are significantly higher and do not support electron transfer.^33,57^ These results indicate that Site-I corresponds to guanine-rich environments that promote electron transfer–mediated quenching, whereas Site-II corresponds to environments where such electron transfer is suppressed. CD measurements were performed to examine whether TMPyP4 binding perturbs the LTR-III GQ structure. The CD spectrum of the LTR-III GQ exhibits characteristic positive bands at ∼265 nm and ∼290 nm, consistent with a hybrid G-quadruplex topology (**Figure 1D**).^13,58^ Notably, these spectral features remain largely unchanged upon addition of TMPyP4, indicating that the overall GQ structure is preserved upon binding (**Figure 1D**). Control experiments with truncated GQ and duplex loop constructs (**Figure S2**) show that the dominant binding interactions arise from the GQ core rather than the duplex hairpin loop. Overall, absorption measurements establish complex formation and stoichiometry, whereas fluorescence titrations reveal two photophysically distinct binding environments, consistent with binding at guanine-rich G-quartet sites and non-guanine regions of the LTR-III GQ.

#### 3.1.2. Time resolved fluorescence studies in buffer

To obtain further insight into the interaction between TMPyP4 and LTR-III GQ, time-resolved fluorescence measurements were carried out using TCSPC. The decay profiles recorded at different GQ concentrations are shown in **Figure 2A**, and the extracted parameters are summarized in **Table 1**. In buffer, in the absence of GQ, TMPyP4 exhibits biexponential decay (**Figure S3, Supporting Information**) with lifetime components of ∼1.5 ns and ∼4.8 ns, where the longer component (∼96%) dominates the emission.^32,59^ The long lifetime component (∼4.8–5 ns) is attributed to monomeric TMPyP4 in aqueous solution and represents the intrinsic excited-state decay in buffer.^32,56,60^ The origin of the shorter component (∼1.5 ns) is less clear and has been reported in the literature with a small contribution (∼3–4%).^32,35^ This component has been attributed to minor populations such as surface-associated species or trace impurities formed during synthesis,^32,35,56^ although no definitive assignment has been established. In the present case, the intensity fraction of this component remains very small and does not significantly influence the overall emission behavior.

**Figure 2.**
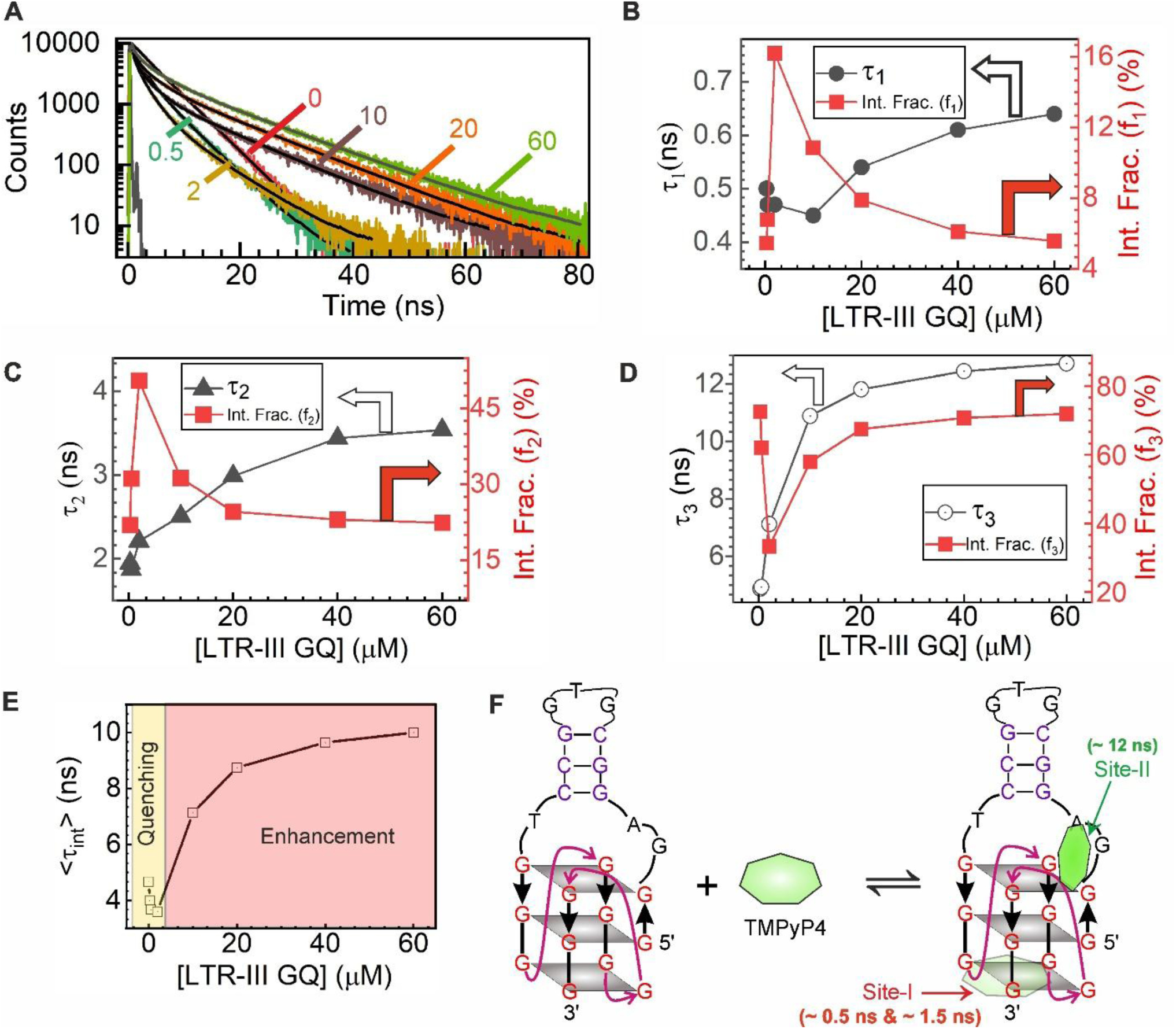
Time-resolved fluorescence studies of TMPyP4 binding with LTR-III GQ in buffer. (A) Fluorescence decay profiles of TMPyP4 (2 μM) recorded at 660 nm upon excitation at 440 nm as a function of increasing GQ concentration. (B–D) Variation of the lifetime components (τ₁, τ₂, τ₃) and their corresponding intensity fractions with GQ concentration, reflecting the evolution of quenching (short lifetime) and non-quenching (long lifetime) populations. (E) Variation of the intensity-weighted average lifetime (⟨τ⟩), showing an initial decrease followed by an increase at higher GQ concentrations, consistent with the bimodal fluorescence behavior observed in steady-state measurements. (F) Schematic representation of the binding modes of TMPyP4 with LTR-III GQ, illustrating quenching at the terminal G-quartet (Site-I) and non-quenching binding at non-guanine regions such as loop and GQ-duplex junction domains (Site-II).

**Table 1:** Fluorescence decay parameters of TMPyP4 obtained in buffer as a function of LTR-III GQ concentrations.

| [LTR-III GQ] ($\mu\text{M}$) | $\tau_1$ (ns) | $\alpha_1$ | $\tau_2$ (ns) | $\alpha_2$ | $\tau_3$ (ns) | $\alpha_3$ | $\langle\tau_{amp}\rangle$ (ns) | $\langle\tau_{int}\rangle$ (ns) |
| --- | --- | --- | --- | --- | --- | --- | --- | --- |
| 0 | $1.52 \pm 0.14$ | 0.11 | $4.79 \pm 0.01$ | 0.89 | | | $4.43 \pm 0.14$ | $4.67 \pm 0.01$ |
| 0.3 | $0.50 \pm 0.03$ | 0.30 | $1.95 \pm 0.18$ | 0.30 | $4.88 \pm 0.02$ | 0.40 | $2.69 \pm 0.19$ | $4.00 \pm 0.02$ |
| 0.5 | $0.47 \pm 0.02$ | 0.33 | $1.87 \pm 0.11$ | 0.38 | $4.94 \pm 0.03$ | 0.29 | $2.30 \pm 0.11$ | $3.68 \pm 0.02$ |
| 2 | $0.47 \pm 0.01$ | 0.56 | $2.21 \pm 0.06$ | 0.37 | $7.13 \pm 0.07$ | 0.08 | $1.62 \pm 0.07$ | $3.60 \pm 0.04$ |
| 10 | $0.45 \pm 0.01$ | 0.57 | $2.51 \pm 0.06$ | 0.30 | $10.89 \pm 0.05$ | 0.13 | $2.40 \pm 0.08$ | $7.14 \pm 0.04$ |
| 20 | $0.54 \pm 0.01$ | 0.51 | $2.99 \pm 0.08$ | 0.29 | $11.82 \pm 0.04$ | 0.20 | $3.49 \pm 0.09$ | $8.75 \pm 0.04$ |
| 40 | $0.62 \pm 0.02$ | 0.44 | $3.44 \pm 0.09$ | 0.30 | $12.45 \pm 0.04$ | 0.25 | $4.48 \pm 0.10$ | $9.65 \pm 0.04$ |
| 60 | $0.64 \pm 0.02$ | 0.42 | $3.55 \pm 0.10$ | 0.31 | $12.72 \pm 0.04$ | 0.27 | $4.84 \pm 0.11$ | $10.00 \pm 0.04$ |

Upon addition of LTR-III GQ, the decay becomes triexponential (**Figure S4, Supporting Information**), indicating the presence of multiple photophysical environments. Three lifetime components are observed (**Table 1** & **Figure 2B–D**): τ₁ (∼0.5–0.6 ns), τ₂ (∼2–3 ns), and τ₃ (∼5–12 ns), which evolve systematically with increasing GQ concentration. To assign these components, control experiments were performed using individual nucleotides at 15 mM concentration (**Figure S5 and Table S1, Supporting Information**). In the presence of GMP, TMPyP4 exhibits two distinct short lifetime components of 0.58 ns and 1.82 ns, closely matching τ₁ and τ₂ observed with GQ, establishing that both originate from guanine-mediated quenching (Site-I). In contrast, AMP and TMP produce dominant long lifetime components of ∼11 ns (**Table S1**), consistent with τ₃ and indicating non-guanine binding environments (Site-II).^31,33,35,57^ The origin of the short lifetime components can therefore be attributed to electron transfer from guanine to the excited state of TMPyP4,^35,57^ as established in steady-state fluorescence measurements. Based on these observations, τ₁ and τ₂ are assigned to TMPyP4 bound at guanine-rich regions of the GQ (Site-I), while τ₃ corresponds to binding at non-guanine regions (Site-II). The presence of two short lifetime components indicates heterogeneity within Site-I, where τ₁ represents a more strongly quenched population and τ₂ a relatively less quenched population within the same binding mode.^33^ However, fluorescence lifetime measurements alone do not provide direct structural information, and therefore no specific assignment of binding geometry or donor–acceptor distance can be made. The evolution of these components with GQ concentration further supports this picture. At low GQ concentrations (0–2 μM), the contributions of τ₁ and τ₂ increase at the expense of τ₃, leading to fluorescence quenching. At higher GQ concentrations (>2 μM), τ₂ shows a gradual increase (Figure 2C), indicating a reduction in quenching efficiency within Site-I. Under these conditions, where GQ is present in excess relative to TMPyP4, it is possible that additional GQ molecules interact with TMPyP4 already bound to a G-quartet, perturbing the local binding environment and reducing quenching efficiency. The long lifetime component (τ₃) arises from binding at Site-II, and its increasing contribution with GQ concentration reflects the progressive population of this binding mode, with the observed lifetime approaching that of the fully bound state (∼12 ns) (**Figure 2D**). As the population shifts from quenched Site-I environments to the less quenched Site-II binding mode, the fluorescence intensity recovers. Consequently, the intensity-weighted average lifetime initially decreases and then increases (**Figure 2E**), consistent with the bimodal fluorescence behavior (**Figure 2F**) observed in steady-state measurements.

#### 3.1.3. Molecular Modelling Studies

The HIV-1 LTR-III target is structurally unique, adopting a hybrid quadruplex-duplex (GQ-D) conformation, with three potential ligand-binding sites: (1) the duplex loop, (2) the bottom G-quartet (G25-G28-G17-G21), and (3) the Q-D junction G-quartet including the loop region (A4, G5, C13).^11^ To investigate the conformational behavior of the system and obtain a stable receptor structure for docking studies, we first evaluated the structural integrity of the apo ion-stabilized LTR-III GQ through a 200 ns MD simulation. The initial trajectory analysis showed a significant increase in the DNA heavy-atom RMSD, reaching ∼7.0 Å between 40 and 80 ns. The large fluctuations in the initial stages correspond to an extensive conformational change, particularly in the flexible loop regions. After 80 ns, the structure reached thermodynamic equilibrium, with the RMSD stabilizing at around 4.5 Å for the rest of the simulation **(Figure 3A)**. Throughout the trajectory, the structural integrity of the central channel was maintained. The average distance between the coordinating guanine O6 atoms and the nearest K^+^ cations was 2.8 Å, while the distance between the two K^+^ cations remained stable at approximately 3.8 Å. Notably, neither of the coordinated K^+^ ions escaped the central channel during the 200 ns simulation **(Figure S6)**, confirming the strong stabilization provided by the explicitly modeled ions.

**Figure 3.**
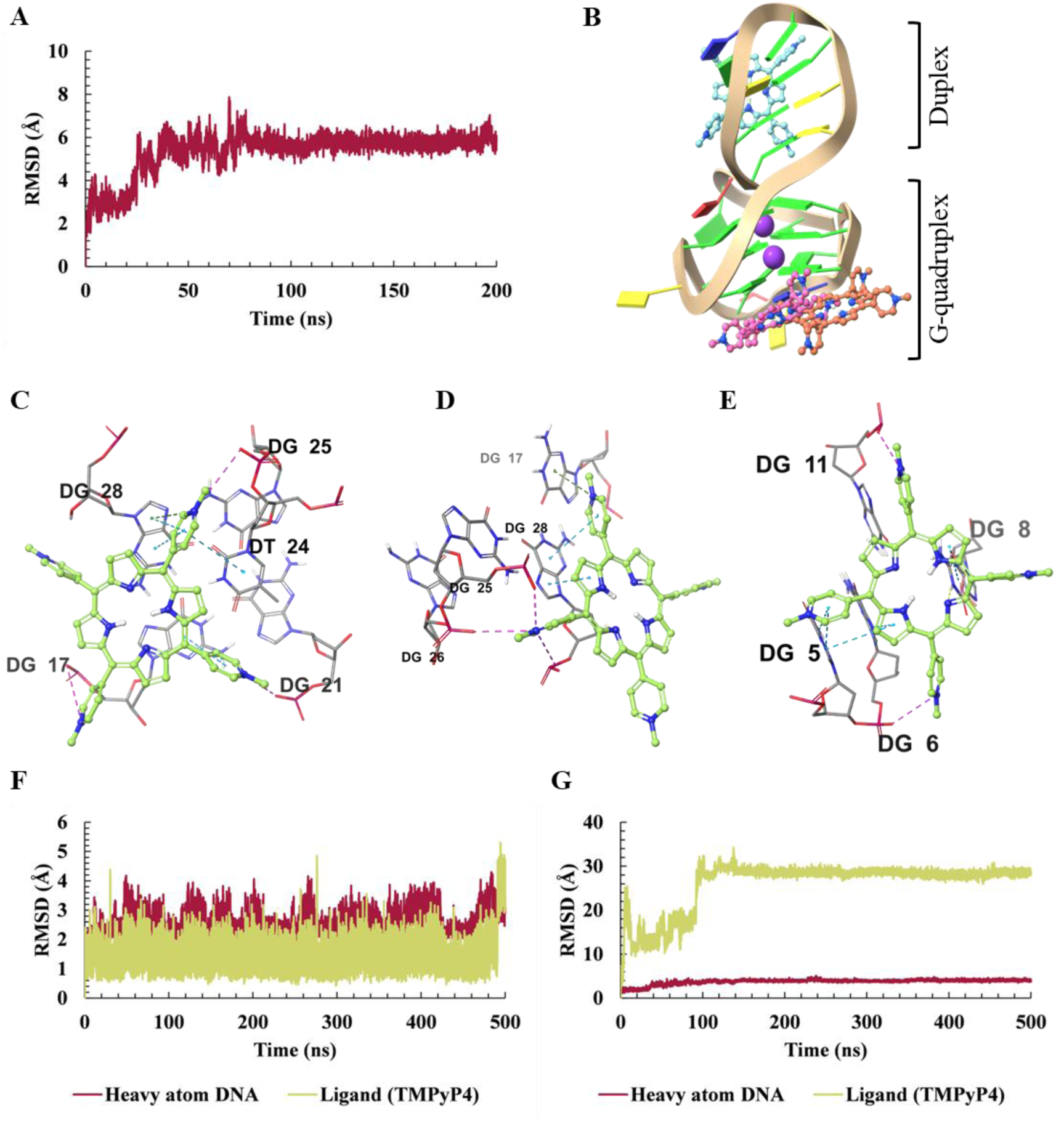
Results of the Molecular docking and MD simulations studies. **(A)** The RMSD of heavy atoms for the apo ion-stabilized LTR-III G-quadruplex over the 200 ns production run, demonstrating initial loop rearrangement and subsequent thermodynamic stabilization. **(B)** Composite structural overview mapping the three evaluated TMPyP4 docked poses onto the LTR-III receptor: the duplex-associated binding site (cyan), the primary bottom G-quartet site (pink), and the secondary GQ associated pose (orange). 3D interaction diagram of the TMPyP4 docked pose localized at the **(C)** bottom G-quartet (primary), **(D)** secondary docked pose derived from the bottom quartet grid in the GQ motif and **(E)** docked pose at the duplex hairpin loop. **(F)** RMSD plot of the ligand (TMPyP4) and the DNA heavy atoms over the 500 ns production run for the primary GQ bottom quartet binding mode. **(G)** RMSD plot for secondary GQ-associated pose over 500 ns, illustrating the initial structural deviation and subsequent stabilization following migration to the GQ-duplex junction.

##### 3.1.3.1 Molecular docking studies to identify possible binding poses

To identify the most plausible binding conformations of TMPyP4, we extracted the most populated cluster from the apo trajectory as a representative receptor. Our experimental observation suggested that the ligand could potentially interact with two distinct binding mode with HIV GQ structures regions; multiple docking grids were generated to cover these structural domains. The Docking poses were screened based on docking scores, MM-GBSA binding free energies, and the extent of π-π stacking interactions with guanine-rich regions. Based on these criteria, three representative binding poses were selected for further analysis: two located within the GQ domain (non-duplex conformations) and one associated with the duplex-bound pose **(Figure 3B)**. The lowest energy GQ bound configuration corresponds to ligand binding at the terminal G-quartet interface, which engages in π-π stacking with G17, G28, and T24 **(Figure 3C)**. A second G-quadruplex associated pose, derived from the same bottom quartet grid, adopts a distinct orientation, characterized by π-π stacking with G28 **(Figure 3D)**. In contrast, the duplex-bound pose engages primarily through stacking with G5, G6, and G8 **(Figure 3E)**. Comparative docking scores and MM-GBSA analysis indicate a clear preference for G-quadruplex binding over the duplex region (**Table 2**).

**Table 2.**
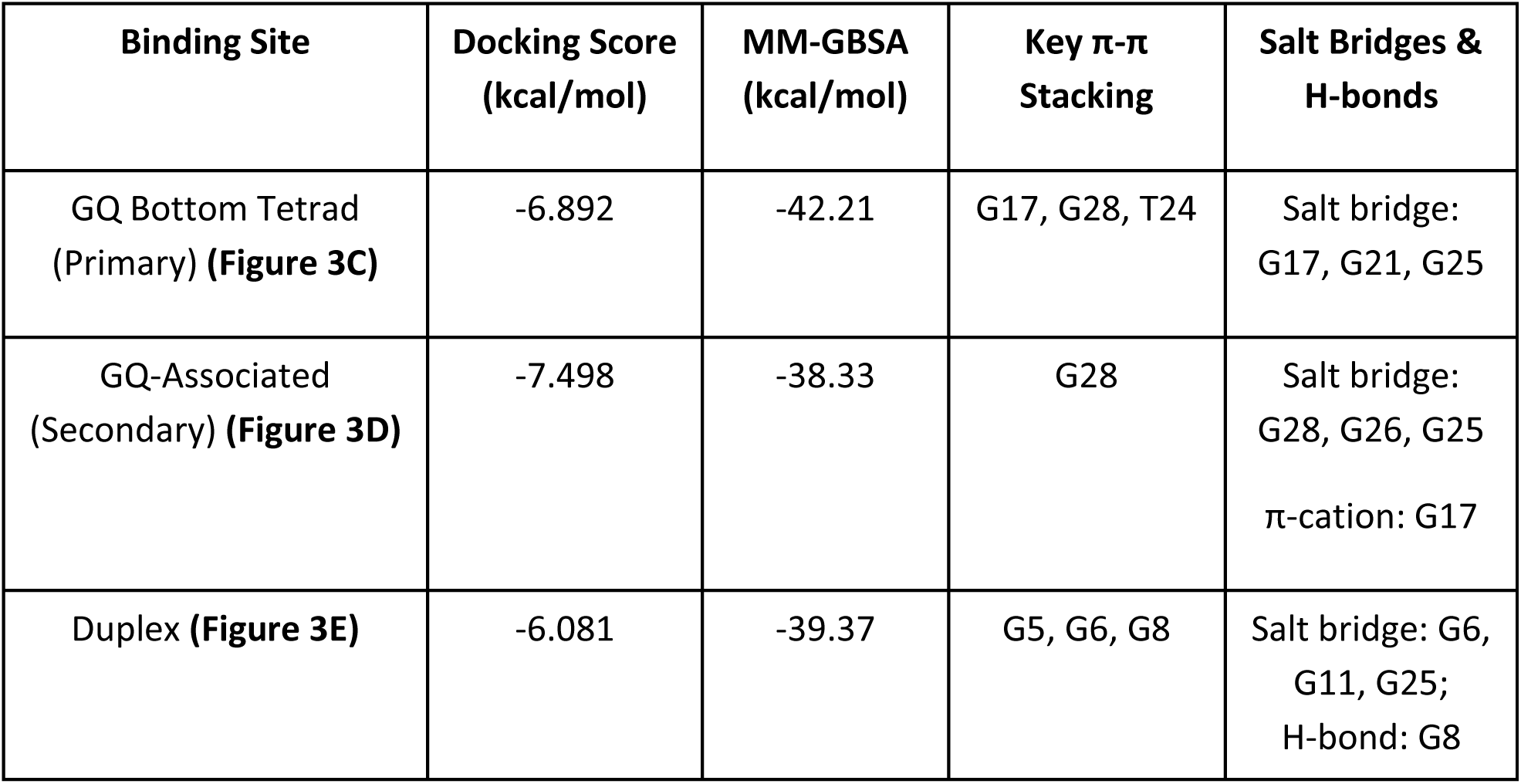
Summary of predicted binding affinities and key residue interactions for TMPyP4 bound to Duplex and GQ regions of HIV-1 LTR-III.

| Binding Site | Docking Score (kcal/mol) | MM-GBSA (kcal/mol) | Key π-π Stacking | Salt Bridges & H-bonds |
| --- | --- | --- | --- | --- |
| GQ Bottom Tetrad (Primary) **(Figure 3C)** | -6.892 | -42.21 | G17, G28, T24 | Salt bridge: G17, G21, G25 |
| GQ-Associated (Secondary) **(Figure 3D)** | -7.498 | -38.33 | G28 | Salt bridge: G28, G26, G25<br>π-cation: G17 |
| Duplex **(Figure 3E)** | -6.081 | -39.37 | G5, G6, G8 | Salt bridge: G6, G11, G25;<br>H-bond: G8 |

##### 3.1.3.2 MD simulations studies to investigate the binding stabilities at different binding poses

To assess the dynamic stability of these binding modes, all three complexes were subjected to 500 ns MD simulations. The RMSD analysis showed that the complex formed at the GQ quartet remained highly stable throughout the trajectory. The Ligand RMSD stabilized around 1.5 Å, while the overall DNA RMSD remained close to 2.6 Å, indicating minimal deviation from the initial binding orientation **(Figure 3F)**. Clustering further confirmed this stability, showing that a single major conformational cluster accounted for about 93% of the simulation. To assess the persistence of the critical stacking interactions at the GQ binding site, we applied center of distance criteria (< 5 Å) and angle criteria (< 45° or > 135°) across the trajectory.^61^ TMPyP4 maintained strong π-π stacking interactions with G28 for about 91% of the simulation and with G17 for around 98%.

In contrast, the alternative GQ-associated pose underwent substantial rearrangement during the early stages of the simulation and subsequently relocated to the GQ-duplex junction region, stabilizing after ∼120ns near residues A4 and A22 **(Figure 3G)**. Clustering revealed a dominant conformational state representing 63% of the trajectory. Interaction analysis revealed sustained contacts with A22 (81%) and A4 (52%). However, the associated binding energetics were less favorable than that of the quartet-bound complex, indicating that this junction represents a secondary binding site. This supports our experimental observation that the secondary binding site (site-II) is dominated by non-guanine bases which is leading to enhanced fluorescence emission and longer lifetime of TMPyP4.

The duplex-bound complex displayed the highest degree of structural fluctuations and failed to maintain a consistent binding orientation throughout the 500 ns trajectory **(Figure S7)**, suggesting weaker binding stability. These trends are aligning with post-MD MM-PBSA binding free energy calculations, which indicated less favorable binding for the duplex-associated complex (−23.50 kcal/mol) compared to the quartet-bound GQ complex (−27.42 kcal/mol) and the GQ-duplex junction associated pose (−25.50 kcal/mol).

Overall, TMPyP4 preferentially interacts with the GQ domains of the HIV-1 LTR-III structure, rather than the duplex hairpin regions. This thermodynamic preference is driven primarily by highly stable intercalative like stacking at the exposed bottom quartet of the GQ, acting in concert with secondary stabilization at the adjacent GQ-duplex junction. Recognizing this dual-site binding behavior offers opportunity for the rational design of ligands. Specifically, these insights strongly advocate for the development of tailored bimodal ligand scaffolds strategically engineered to combine both GQ bottom-quartet and GQ-duplex binding moieties.

### 3.2. Mechanistic insights into the binding in BSA droplets formed in BSA-PEG ATPS

#### 3.2.1. Formation and microscopic characterization of the BSA-PEG ATPS

To investigate the binding interaction in a condensate-like environment, a BSA-PEG ATPS was employed. PEG acts as a macromolecular crowder that enhances interactions between BSA molecules through excluded volume effects and reduced solvent availability, thereby driving phase separation into a protein-rich condensed phase.^37^ The phase behavior was examined by turbidometric measurements by varying the relative ratio of BSA and PEG at a fixed total concentration of 240 mg/mL. The absorbance at 400 nm is used as a measure of turbidity and arises primarily from light scattering by phase-separated domains.^17^ The neat PEG solution (240 mg/mL) shows a relatively high apparent absorbance despite being optically clear, which can be attributed to increased light scattering from the highly viscous PEG solution. Upon increasing the BSA/PEG ratio, the absorbance remains nearly constant up to a ratio of 0.25, indicating a single-phase system, followed by a sharp increase beyond ∼0.3, consistent with the onset of phase separation due to the formation of scattering droplets (**Figure 4A,B**). At higher ratios (>0.9), the turbidity decreases, suggesting a return to a homogeneous phase. Phase contrast imaging at 70 mg/mL BSA and 170 mg/mL PEG, corresponding to the region of maximum turbidity, reveals micron-sized spherical droplets (**Figure 4C**), confirming LLPS. To verify the composition of these droplets, a small fraction of Alexa 647-labeled BSA (1 mg/mL) was introduced into the same system. The fluorescence signal is localized within the droplets and overlaps with the DIC features (**Figure 4D**), indicating that the condensates are enriched in BSA.

**Figure 4:**
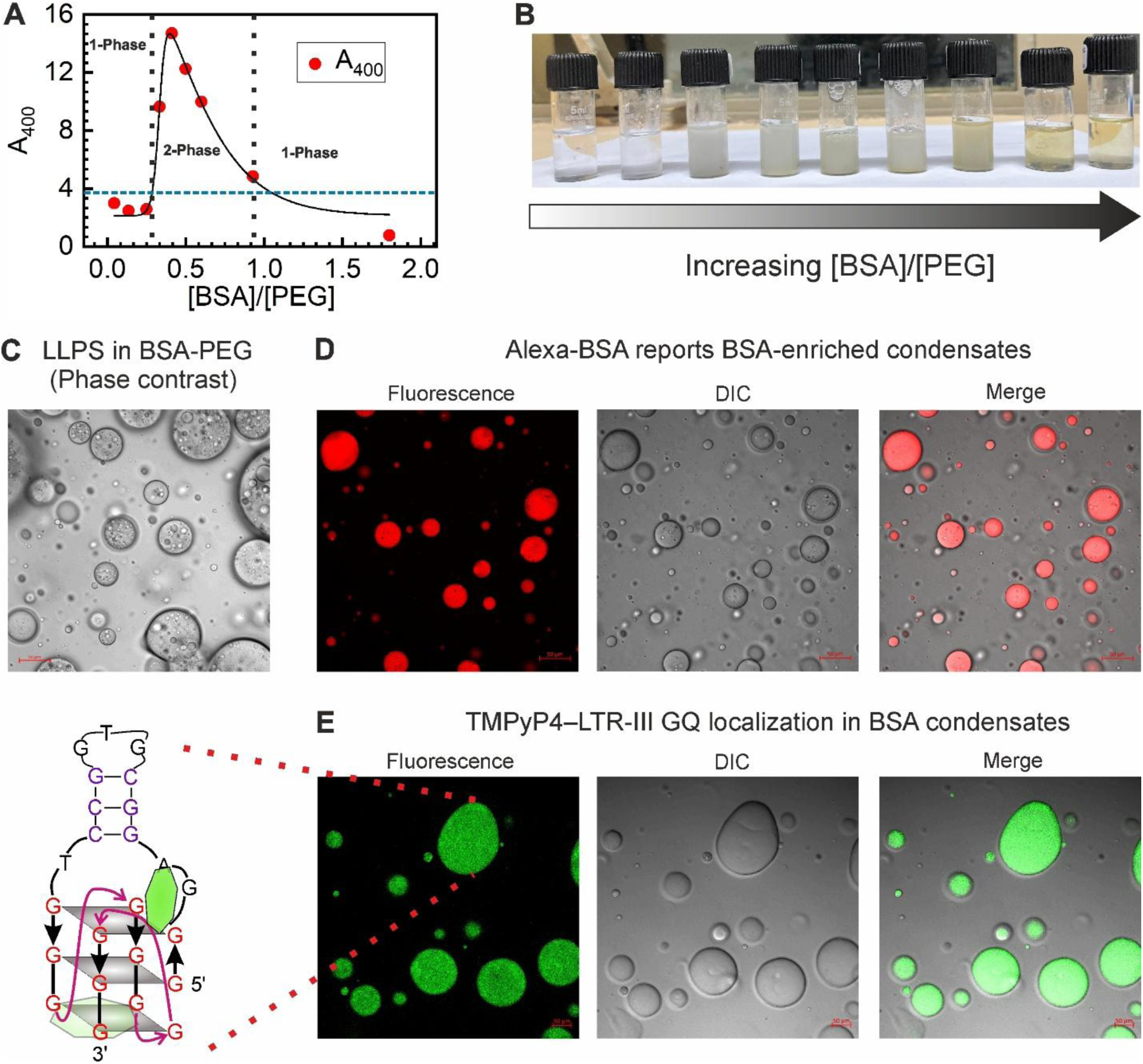
Formation and microscopic characterization of BSA–PEG condensates, and accumulation of TMPyP4–GQ within the droplet phase. (A) Absorbance at 400 nm (A₄₀₀) as a function of [BSA]/[PEG], indicating a transition from a single-phase to a two-phase regime. (B) Photographs of BSA–PEG mixtures showing macroscopic phase separation. (C) Phase contrast image demonstrating liquid–liquid phase separation (LLPS) and formation of micron-sized BSA condensates. (D) Fluorescence, DIC, and merged images of condensates probed with a small fraction of Alexa-labeled BSA mixed with unlabeled BSA in the presence of PEG; fluorescence co-localizes with the droplets, indicating that the condensed phase is enriched in BSA. (E) Fluorescence, DIC, and merged images of BSA–PEG condensates containing TMPyP4 and LTR-III G-quadruplex DNA, showing accumulation of TMPyP4 fluorescence within the droplet phase. **Scale bars: 50 µm (all panels).**

Having established that the condensates are BSA-rich, TMPyP4 and LTR-III G-quadruplex (GQ) DNA were introduced into the same BSA–PEG system to examine their distribution within the droplets. The mixture was prepared using 30 μM GQ DNA and 2 μM TMPyP4, where TMPyP4 exhibits enhanced fluorescence upon binding to GQ DNA. The fluorescence and DIC images (**Figure 4E**) show that TMPyP4 fluorescence is strongly localized within the droplets and co-localizes with the DIC features, indicating preferential accumulation in the BSA-rich phase. The fluorescence intensity within the droplets is significantly higher than that of the surrounding medium. At these concentrations, TMPyP4 is expected to be predominantly bound to GQ DNA; thus, its localization within the droplets suggests partitioning of the TMPyP4–GQ complex into the condensed phase. This behavior is consistent with PEG-induced excluded volume effects, which disfavour the presence of macromolecular species in the PEG-rich phase and consequently promote their accumulation within the BSA-rich droplets.

#### 3.2.2. Binding study between TMPyP4 and LTR-III GQ inside the BSA droplet using steady state fluorescence

Fluorescence titrations were performed in BSA–PEG phase-separated droplets to examine how a condensate-like environment influences ligand–GQ recognition under conditions identical to those used in buffer. As established in the structural characterization section, TMPyP4 preferentially partitions into the BSA-rich phase (**Figure 5A**). In the absence of GQ, the emission spectrum of TMPyP4 within the condensate is clearly resolved into two bands at ∼653 nm and ∼714 nm (**Figure 5B**), in contrast to the broad, unresolved emission observed in buffer. This spectral resolution arises from binding of TMPyP4 to hydrophobic subdomains of BSA, where reduced local polarity and restricted conformational flexibility diminish coupling between the locally excited (S₁) and charge-transfer (CT) states, resulting in well-defined emission features.^31,56^ Upon addition of LTR-III GQ, a bimodal fluorescence response is observed (**Figure 5B,C**), characterized by an initial decrease in emission intensity up to ∼10 μM followed by recovery and eventual saturation at higher GQ concentrations. This behavior closely mirrors that observed in buffer, indicating that the two distinct binding modes—guanine-mediated quenching (Site-I) and and a non-guanine binding mode associated with longer lifetime (Site-II)—remain operative within the condensate phase. Both emission bands exhibit a systematic red shift with increasing GQ concentration (653 → 661 nm and 713 → 724 nm), reflecting changes in the local environment of TMPyP4 as it redistributes from protein-associated sites to GQ-bound states. Binding isotherms constructed from the quenching and enhancement regimes (**Figure 5D,E**) yield dissociation constants of 530 ± 70 nM (Site-I) and 10.1 ± 1.5 μM (Site-II), respectively. These values are modestly higher than those obtained in buffer (408.7 ± 50.0 nM and 6.7 ± 0.7 μM; **Figure 5F**). The increase in dissociation constants in the condensate phase indicates a slight shift in equilibrium toward the unbound state. This shift is more clearly visualized in terms of the dissociation free energy (ΔG^0^_diss_ = −RT ln K_d_). As shown in **Figure 5G**, ΔG^0^_diss_ becomes slightly less positive in the condensate phase for both binding modes, with values changing from 8.71 ± 0.07 to 8.56 ± 0.08 kcal mol⁻¹ for Site-I and from 7.05 ± 0.06 to 6.81 ± 0.09 kcal mol⁻¹ for Site-II, corresponding to ΔΔG^0^_diss_ ≈ −0.15 ± 0.11 kcal mol⁻¹ (Site-I) and −0.24 ± 0.11 kcal mol⁻¹ (Site-II) (**Figure 5H**). The magnitude of this change is very small (<0.3 kcal mol⁻¹), indicating that dissociation is only weakly favored in the condensate environment despite the substantial change in medium.

**Figure 5:**
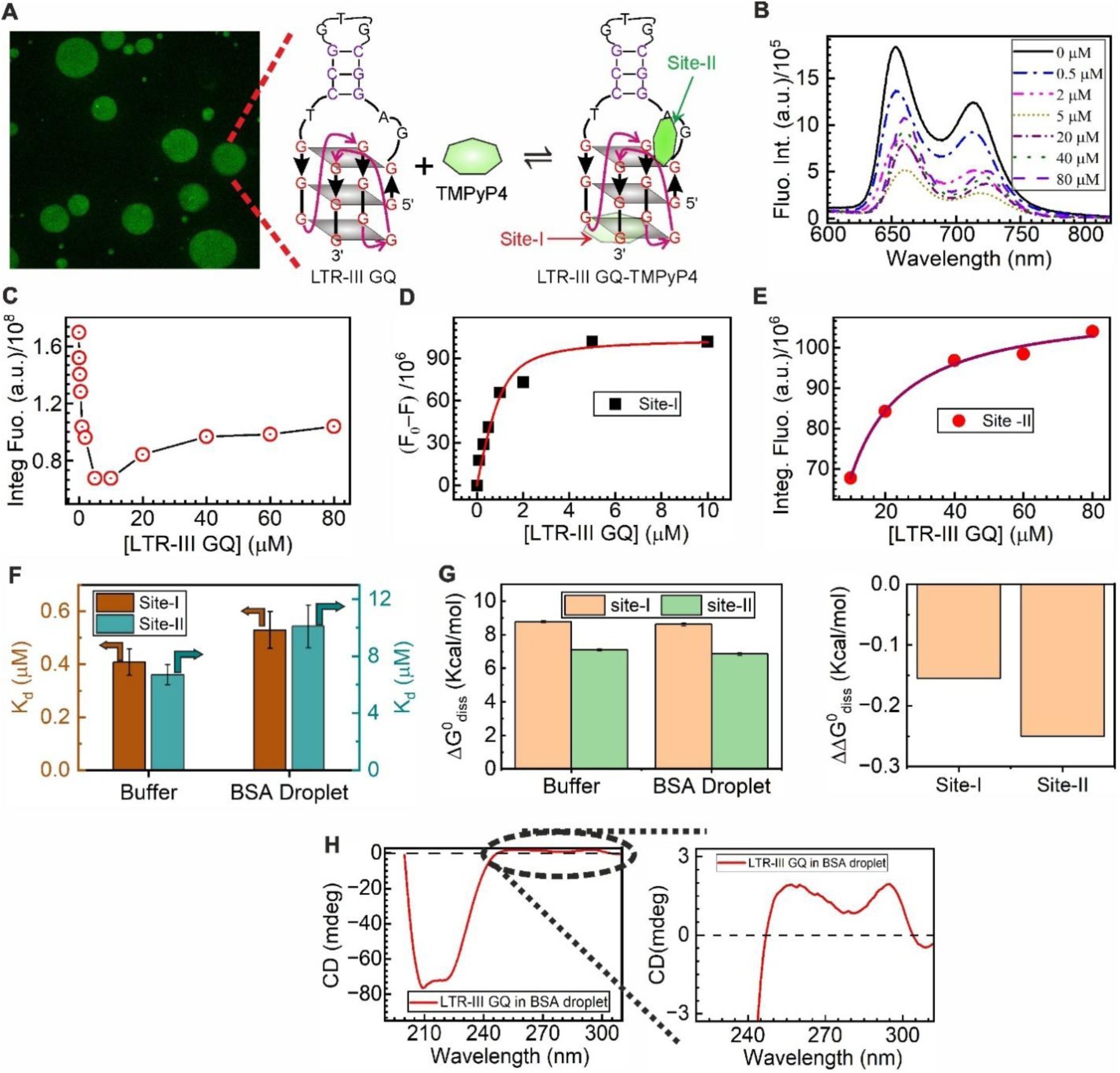
Results of the Binding study in BSA–PEG phase-separated condensates. (A) Fluorescence microscopy images showing preferential partitioning of both TMPyP4 and LTR-III GQ into the BSA-rich droplet phase. (B) Steady-state fluorescence spectra of TMPyP4 (2 μM) in condensates upon titration with LTR-III GQ (0–80 μM), displaying two well-resolved emission bands. (C) Corresponding variation of fluorescence intensity showing a bimodal response with initial quenching followed by recovery at higher GQ concentrations. (D,E) Binding isotherms for Site-I (quenching-dominated) and Site-II (emission-recovery regime), respectively, obtained from integrated fluorescence intensity and fitted using the model described in the Methods section. (F) Comparison of dissociation constants (K_d_) for Site-I and Site-II in buffer and condensate, indicating a modest increase in K_d_ in the BSA-rich phase. (G) Dissociation free energy (ΔG^0^_diss_) and corresponding change in free energy (ΔΔG^0^_diss_) for both sites, showing only a small perturbation in binding energetics (<0.3 kcal mol⁻¹) upon transfer to the condensate. (H) Circular dichroism spectra of LTR-III GQ in buffer and condensates confirming retention of the hybrid GQ topology along with the characteristic α-helical signature of BSA.

Circular dichroism measurements confirm that the LTR-III G-quadruplex retains its hybrid topology within the condensate, as evidenced by the characteristic positive bands at ∼260 nm and ∼295 nm (**Figure 5H**), while the secondary structure of BSA remains preserved.^19,38^ Thus, neither the GQ structure nor the nature of the binding sites is disrupted in the droplet phase. The small change in dissociation free energy is noteworthy, as molecular crowding is known to significantly influence ligand–nucleic acid interactions through coupled effects on hydration, solvent environment, and molecular mobility.^62–65^ In the case of TMPyP4, binding to G-quadruplex DNA has been reported to involve uptake of water molecules,^62–64^ suggesting that reduced water activity in the condensate phase would thermodynamically disfavor complex formation. In addition, the high viscosity of the BSA-rich phase can slow down molecular diffusion, reducing the frequency of productive ligand–GQ encounters and thereby contributing to a slight shift toward the unbound state.^63,66^ However, these destabilizing contributions are counterbalanced by excluded volume effects that increase the effective concentration of interacting species within the crowded environment.^21,27,67,68^ The resulting small net change in ΔG^0^_diss_ therefore reflects a near-compensation between these opposing contributions, indicating that the overall energetic landscape governing TMPyP4–GQ interaction remains largely preserved. Importantly, the retention of the bimodal fluorescence signature demonstrates that not only the affinity but also the binding topology remains intact, highlighting that ligand recognition of the HIV-1 LTR-III G-quadruplex is remarkably robust within condensate-like environments.

#### 3.2.3. Time resolved fluorescence study inside BSA rich condensates

To examine how the condensate environment influences the interaction of TMPyP4 with LTR-III GQ, time-resolved fluorescence measurements were carried out under identical conditions (**Figure 6A**, **Table 3**). In BSA-rich condensates, even in the absence of GQ, the fluorescence decay of TMPyP4 is triexponential (**Figure S8**), with lifetime components of τ₁ = 0.11 ns, τ₂ = 1.60 ns, and τ₃ = 8.78 ns. Although the shortest component dominates in population, its contribution to the overall emission is small due to strong quenching, and the emission is primarily governed by the long lifetime component (**Figure 6B–D**). Notably, this long lifetime component (∼8.78 ns) is significantly longer than that observed in buffer (∼5 ns), indicating suppression of nonradiative decay pathways within the viscous and dense protein environment. The short lifetime components are attributed to strongly quenched populations arising from interactions of TMPyP4 with BSA, likely involving electron transfer from nearby amino acid residues to excited TMPyP4. In contrast, the long lifetime component reflects TMPyP4 in an environment where nonradiative decay pathways are suppressed due to reduced molecular mobility and conformational flexibility within the condensate.

**Figure 6.**
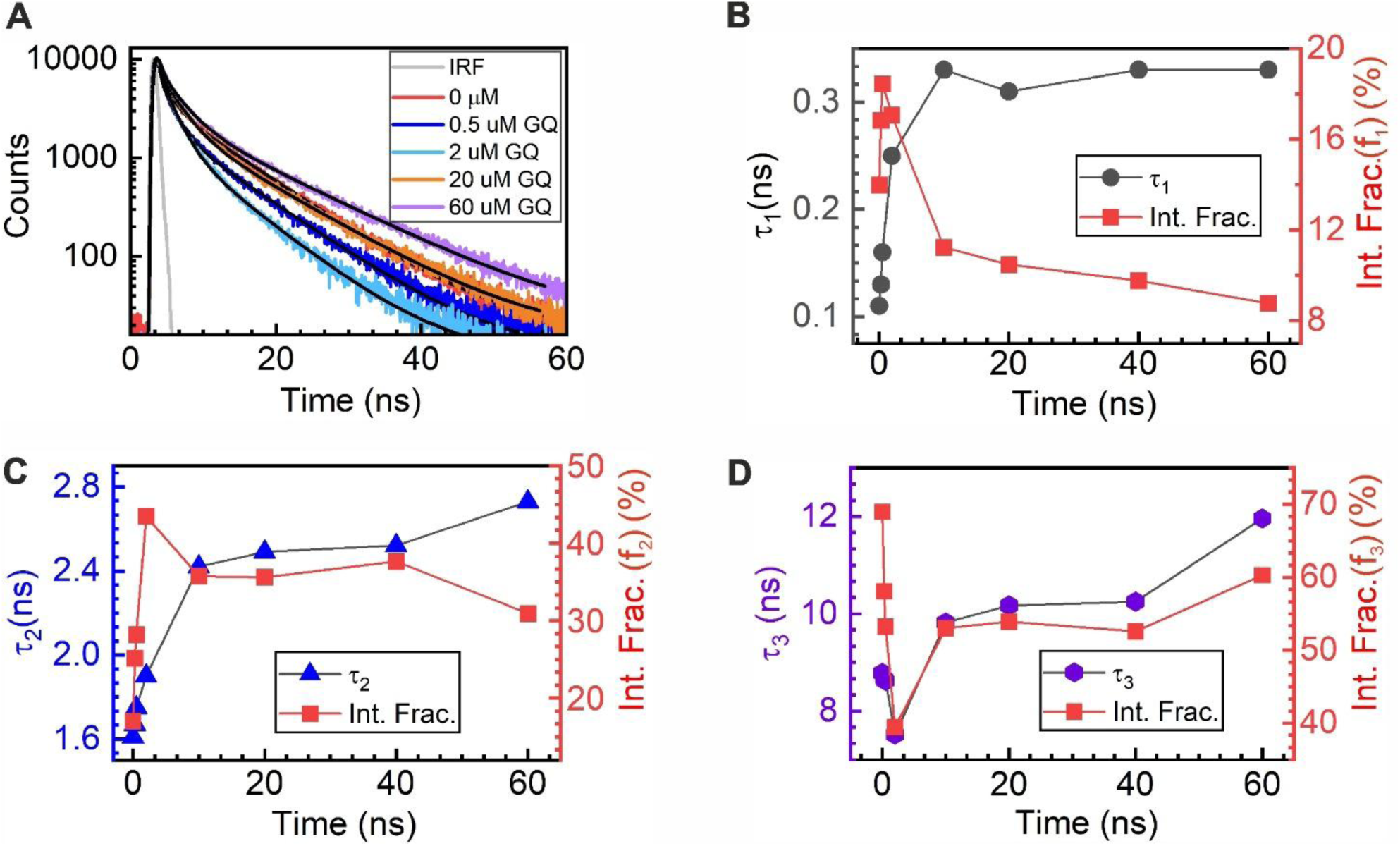
TCSPC studies of TMPyP4 (2 μM) in BSA-rich condensates at varying LTR-III GQ concentrations. (A) Representative fluorescence decay profiles of TMPyP4 in the absence and presence of GQ. (B–D) Variation of lifetime components (τ₁, τ₂, τ₃) and their corresponding intensity fractions with increasing GQ concentration.

**Table 3:** Time-resolved fluorescence decay parameters of TMPyP4 (2 μM) in BSA-rich condensates at varying concentrations of LTR-III GQ.

| [LTR-III GQ] ($\mu\text{M}$) | $\tau_1$ (ns) | $\alpha_1$ | $\tau_2$ (ns) | $\alpha_2$ | $\tau_3$ (ns) | $\alpha_3$ | $\langle\tau\rangle_{amp}$ (ns) | $\langle\tau\rangle_{int}$ (ns) |
| --- | --- | --- | --- | --- | --- | --- | --- | --- |
| 0 | $0.11 \pm 0.007$ | 0.87 | $1.60 \pm 0.03$ | 0.07 | $8.78 \pm 0.03$ | 0.06 | $0.73 \pm 0.04$ | $6.34 \pm 0.04$ |
| 0.3 | $0.13 \pm 0.003$ | 0.85 | $1.67 \pm 0.03$ | 0.10 | $8.67 \pm 0.03$ | 0.05 | $0.71 \pm 0.04$ | $5.47 \pm 0.03$ |
| 0.5 | $0.16 \pm 0.006$ | 0.84 | $1.75 \pm 0.03$ | 0.12 | $8.62 \pm 0.04$ | 0.04 | $0.69 \pm 0.05$ | $5.11 \pm 0.02$ |
| 2 | $0.25 \pm 0.01$ | 0.71 | $1.90 \pm 0.03$ | 0.24 | $7.52 \pm 0.05$ | 0.05 | $1.01 \pm 0.06$ | $3.84 \pm 0.03$ |
| 10 | $0.33 \pm 0.01$ | 0.62 | $2.42 \pm 0.05$ | 0.28 | $9.82 \pm 0.05$ | 0.10 | $1.86 \pm 0.07$ | $6.11 \pm 0.05$ |
| 20 | $0.31 \pm 0.01$ | 0.63 | $2.49 \pm 0.05$ | 0.27 | $10.17 \pm 0.05$ | 0.10 | $1.88 \pm 0.07$ | $6.40 \pm 0.05$ |
| 40 | $0.33 \pm 0.01$ | 0.60 | $2.52 \pm 0.04$ | 0.30 | $10.25 \pm 0.05$ | 0.10 | $1.98 \pm 0.06$ | $6.37 \pm 0.04$ |
| 60 | $0.33 \pm 0.01$ | 0.62 | $2.73 \pm 0.10$ | 0.26 | $11.96 \pm 0.05$ | 0.12 | $2.35 \pm 0.08$ | $8.08 \pm 0.05$ |

Upon addition of LTR-III GQ, the decay remains triexponential (**Figure S9**), and the evolution of the lifetime components follows trends similar to those observed in buffer (**Figure 6B–D**). Both τ₁ and τ₂ remain nearly unchanged at low GQ concentrations and begin to increase at higher concentrations (**Table 3** & **Figure 6B–C**). Beyond ∼10 μM GQ, τ₁ shows little further change and remains nearly constant, whereas τ₂ shows a slight further increase before reaching a plateau at higher concentrations. At low GQ concentrations (0–2 μM), the intensity fractions of the short lifetime components increase, accompanied by a decrease in the long lifetime component, leading to fluorescence quenching (**Figure 6B-D**). At higher GQ concentrations (> 2 μM), the contribution of the long lifetime component (τ₃) increases, while that of the shortest lifetime component (τ_1_) decreases (**Figure 6B&D**), leading to recovery of fluorescence intensity. Similar to the behavior observed in buffer, the short lifetime components (τ₁ and τ₂) inside the condensates are attributed to quenched TMPyP4 populations associated with guanine-rich binding environments, where electron transfer from guanine to excited TMPyP4 remains operative. In contrast, the long lifetime component (τ₃) corresponds to binding environments where such quenching is suppressed, analogous to the non-guanine Site-II identified in buffer. The initial increase in the intensity fractions of the short lifetime components at low GQ concentrations (**Figure 6A&B**) therefore reflects preferential population of guanine-rich binding sites, leading to fluorescence quenching. At higher GQ concentrations, the increasing contribution of τ₃ together with the reduction in the short lifetime populations indicates increasing population of non-guanine binding environments, resulting in fluorescence recovery. Consequently, the intensity-weighted average lifetime initially decreases and then increases (**Table 3**), consistent with steady-state observations.

Overall, despite the additional complexity introduced by the BSA-rich environment, the key features of TMPyP4 interaction with LTR-III GQ are preserved. The presence of multiple lifetime components associated with distinct binding environments, along with their systematic evolution with GQ concentration, indicates that the two binding modes identified in buffer remain operative within the condensate phase.

## Conclusions

In this work, we elucidate the molecular basis of TMPyP4 recognition of the HIV-1 LTR-III GQ under both dilute conditions and in BSA protein condensates. Steady-state and time-resolved fluorescence measurements reveal a dual binding behavior that is not discernible from absorption spectroscopy, arising from two distinct binding environments within the LTR-III GQ scaffold. A high-affinity binding mode associated with guanine-rich regions (Site-I) leads to efficient fluorescence quenching, whereas a weaker binding mode associated with non-guanine regions (Site-II) gives rise to enhanced and long-lived emission. Control experiments with individual nucleotides establish that quenching originates from electron transfer from ground-state guanine to excited TMPyP4, thereby identifying Site-I as a guanine-rich binding environment.

Molecular docking and MD simulations further support the existence of multiple binding modes. The simulations reveal that TMPyP4 preferentially binds at the terminal G-quartet through stable π–π stacking interactions, while a secondary binding mode is stabilized near the quadruplex–duplex junction. In contrast, duplex-associated binding exhibits substantially lower stability during MD simulations, consistent with the experimental observation that the dominant interaction originates from the GQ domain rather than the duplex loop. Together, the spectroscopic and computational results indicate that the primary high-affinity binding mode arises from interactions with guanine-rich G-quartet regions, while the weaker binding mode is associated with non-guanine environments located near the loop and Q-D junction regions.

Time-resolved fluorescence measurements provide direct insight into the population dynamics of these binding modes. The systematic evolution of the lifetime components and their corresponding intensity fractions with increasing GQ concentration reveals a progressive shift in population from Site-I to Site-II, accounting for the observed bimodal fluorescence response. The presence of multiple short lifetime components within Site-I further indicates heterogeneity within the guanine-mediated quenching environment.

Importantly, both TMPyP4 and LTR-III GQ preferentially partition into BSA-rich condensates formed in the BSA–PEG aqueous two-phase system, while the hybrid G-quadruplex structure remains preserved inside the condensates. Despite the crowded, confined, and viscous condensate environment, the dual binding modes and their associated excited-state signatures remain intact. Only a slight reduction in binding affinity is observed inside the condensates. This small change likely arises from near-compensating contributions of opposing physicochemical effects, where reduced water activity and restricted molecular mobility slightly weaken binding, while excluded volume effects favor ligand–GQ association. As a result, the overall energetic landscape governing TMPyP4 recognition remains largely preserved.

Overall, this work demonstrates that both the structure and ligand recognition properties of the HIV-1 LTR-III GQ remain preserved in condensate-like environments. More broadly, these findings establish fluorescence spectroscopy as a powerful approach for resolving hidden binding heterogeneity in GQ–ligand interactions and provide molecular-level insight into ligand recognition of viral GQs under biologically relevant protein condensate conditions.

## Supporting information

Data is enclosed as supplementary files

## Acknowledgement

This work is supported by the Start-up Research Grant (Project: SERB/SRG/2022/000341) from the Science and Engineering Research Board (SERB), now known as the Anusandhan National Research Foundation (ANRF), Government of India. We also acknowledge financial support from the Research Initiation Grant (RIG) and Additional Competitive Research Grant (ACRG) of BITS Pilani. S. Pradhan, S. M. T., S. Sharma, and A. P. S. acknowledge Institute Fellowships from BITS Pilani. S. Sharma also acknowledges SERB (now ANRF) for providing a JRF fellowship from November 2022 to September 2024. We thank Dr. Sonali Gangwar at the Central Research Facility (CRF), IIT Delhi, for assistance with the CD measurements. We also thank Sophisticated Analytical and Technical Help Institute (SATHI) at IIT Delhi for providing access to use the time resolved fluorescence facility.

