## Supplementary material for "Mechanistic Insights into TMPyP4 Recognition of the HIV-1 LTR-III G-Quadruplex in Dilute and Protein Condensate Environments Reveal Hidden Dual Binding Modes": Data is enclosed as supplementary files

### Contents:

1. Binding Model and Fitting Procedure

2. Results of the steady state Fluorescence titration measurements to study the binding between TMPyP4 with individual mononucleotides. (**Figure S1**)

3. Binding of TMPyP4 to truncated LTR-III G-quadruplex (GQ) and duplex loop constructs. (**Figure S2**)

4. Comparison of monoexponential and biexponential fits to the fluorescence decay of TMPyP4 in buffer in the absence of LTR-III G-quadruplex (0 μM GQ). (**Figure S3**)

5. Comparison of monoexponential, biexponential, and triexponential fits to the fluorescence decay profiles of TMPyP4 in the presence of LTR-III G-quadruplex (GQ) in buffer. (**Figure S4**)

6. Time-resolved fluorescence decay profiles of TMPyP4 in the presence of different nucleotides at varying concentrations. (**Figure S5**)

7. TCSPC decay parameters of TMPyP4 (2 μM) in the absence and presence of mononucleotides (GMP, AMP, TMP, and CMP). (**Table S1**)

8. MD simulation run of the ion induced stabilization of LTR-III GQ. (**Figure S6**)

9. Study of the binding stability of the TMPyP4-LTR-III GQ complex for binding pose at the duplex domain. (**Figure S7**)

10. Comparison of (A) monoexponential, (B) biexponential, and (C) triexponential fits to the fluorescence decay profiles of TMPyP4 (2 μM) in BSA-PEG condensates. (**Figure S8**)

11. Comparison of (A) monoexponential, (B) biexponential, and (C) triexponential fits to the fluorescence decay profiles of TMPyP4 (2 μM) in the presence of 60 μM LTR-III GQ in BSA-PEG condensates. (**Figure S9**)

**1. Binding Model and Fitting Procedure**:

Binding curves were analysed using an **independent and equivalent binding sites model**. The LTR-III G-quadruplex (GQ) was assumed to contain *n* identical, non-interacting binding sites for TMPyP4. The overall binding equilibrium may be written as

$$GQ + nTMPyP4 \rightleftharpoons GQ - (TMPyP4)_n$$

The microscopic binding process corresponds to the binding of a single TMPyP4 molecule to one site on the GQ:

$$GQ + TMPyP4 \rightleftharpoons GQ - TMPyP4$$

The microscopic dissociation constant ($K_d$) for one binding site is defined as

$$K_d = \frac{[Free\ binding\ sites][TMPyP4]_f}{[GQ - TMPyP4]_b} \qquad \text{(S1)}$$

where $[\text{TMPyP4}]_f$is the concentration of free ligand and $[\text{GQ–TMPyP4}]_b$is the concentration of bound ligand. Let the total concentration of GQ be $[\text{GQ}]_0 = x$. Since each GQ molecule contains *n* equivalent binding sites, the total concentration of binding sites is $nx$. If the concentration of bound ligand is $B$, then the mass-balance relations are

$$[\text{TMPyP4}]_f = L_0 - B \text{ and } [\text{free binding sites}] = nx - B$$

where $L_0$is the total TMPyP4 concentration. Substituting these expressions into **Eq. S1** gives

$$K_d = \frac{(nx - B)(L_0 - B)}{B}$$

which can be rearranged to yield the quadratic equation in B

$$B^2 - (nx + L_0 + K_d)B + nxL_0 = 0 \qquad \text{(S2)}$$

The larger root is not physically acceptable as it would correspond to a bound ligand concentration exceeding either the total ligand concentration or the total number of available binding sites. The experimentally observed fluorescence response varies linearly with the concentration of bound TMPyP4. The fluorescence intensity measured in the presence and absence of GQ are denoted by $F$and $F_0$, respectively. For **fluorescence quenching experiments**, the magnitude of quenching is proportional to the concentration of bound TMPyP4 and is described by

$$F_0 - F = \alpha \frac{(nx + L_0 + K_d) - \sqrt{(nx + L_0 + K_d)^2 - 4nxL_0}}{2} \qquad \text{(S4)}$$

where $\alpha$is the proportionality constant relating the fluorescence response to the concentration of bound TMPyP4. For **fluorescence enhancement experiments**, the fluorescence increase upon binding is given by

$$F - F'_0 = \alpha \frac{(nx + L_0 + K_d) - \sqrt{(nx + L_0 + K_d)^2 - 4nxL_0}}{2} \qquad \text{(S5)}$$

where $F$represents the fluorescence intensity at different GQ concentrations in the fluorescence-enhancement regime, and $F_0'$corresponds to the minimum fluorescence intensity obtained during the titration, i.e., the fluorescence intensity immediately before the onset of fluorescence enhancement. All the nonlinear least-squares fitting was performed using **OriginPro 2023** employing the **Levenberg–Marquardt algorithm**.

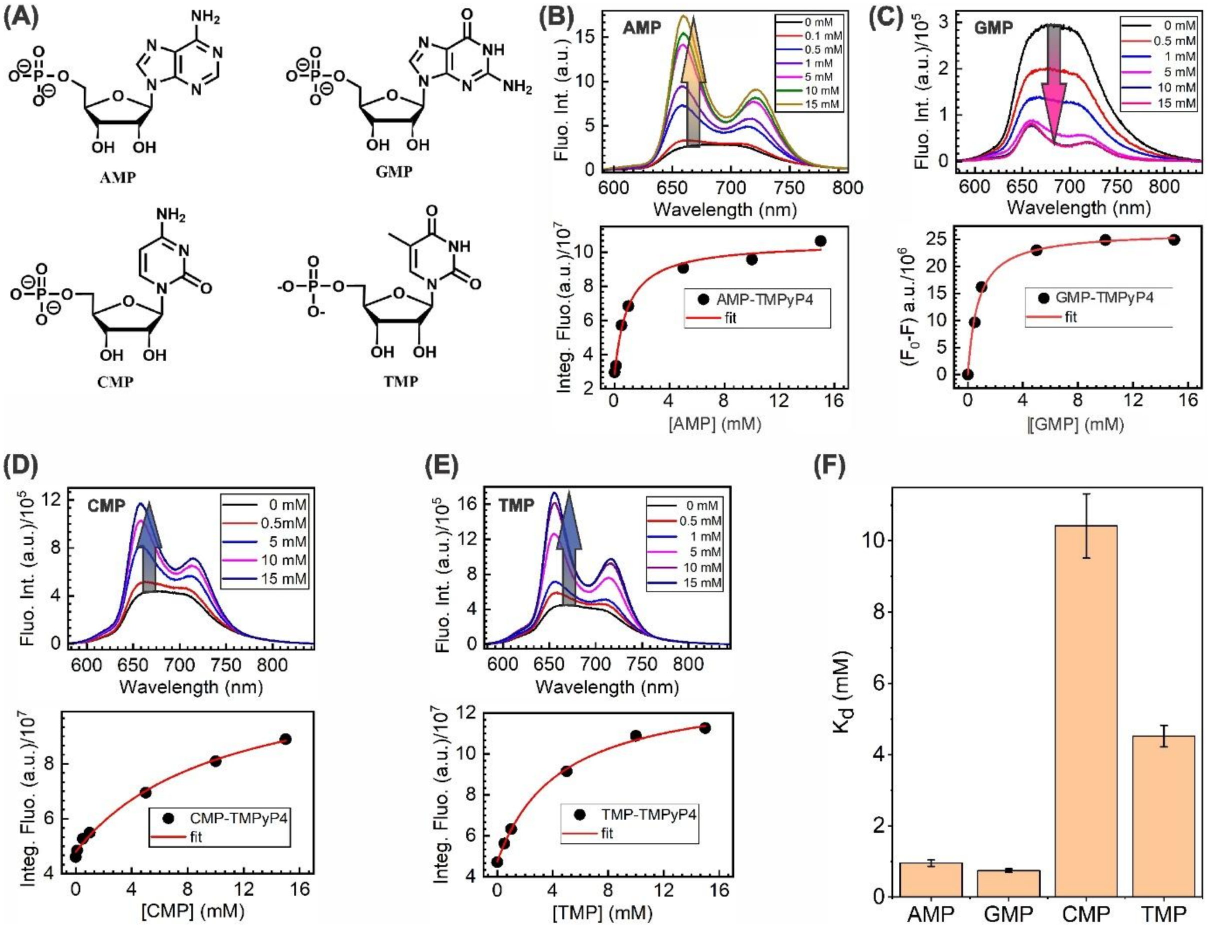


**Figure S1. Fluorescence response of TMPyP4 (2 μM) in the presence of individual nucleotides.** (A) Chemical structures of AMP, GMP, CMP, and TMP. (B–E) Steady-state fluorescence spectra of TMPyP4 upon titration with increasing concentrations of AMP, GMP, CMP, and TMP, respectively, along with the corresponding integrated fluorescence intensity plots as a function of nucleotide concentration. (F) Apparent dissociation constants ($K_d$) obtained from fitting the integrated fluorescence data for each nucleotide.

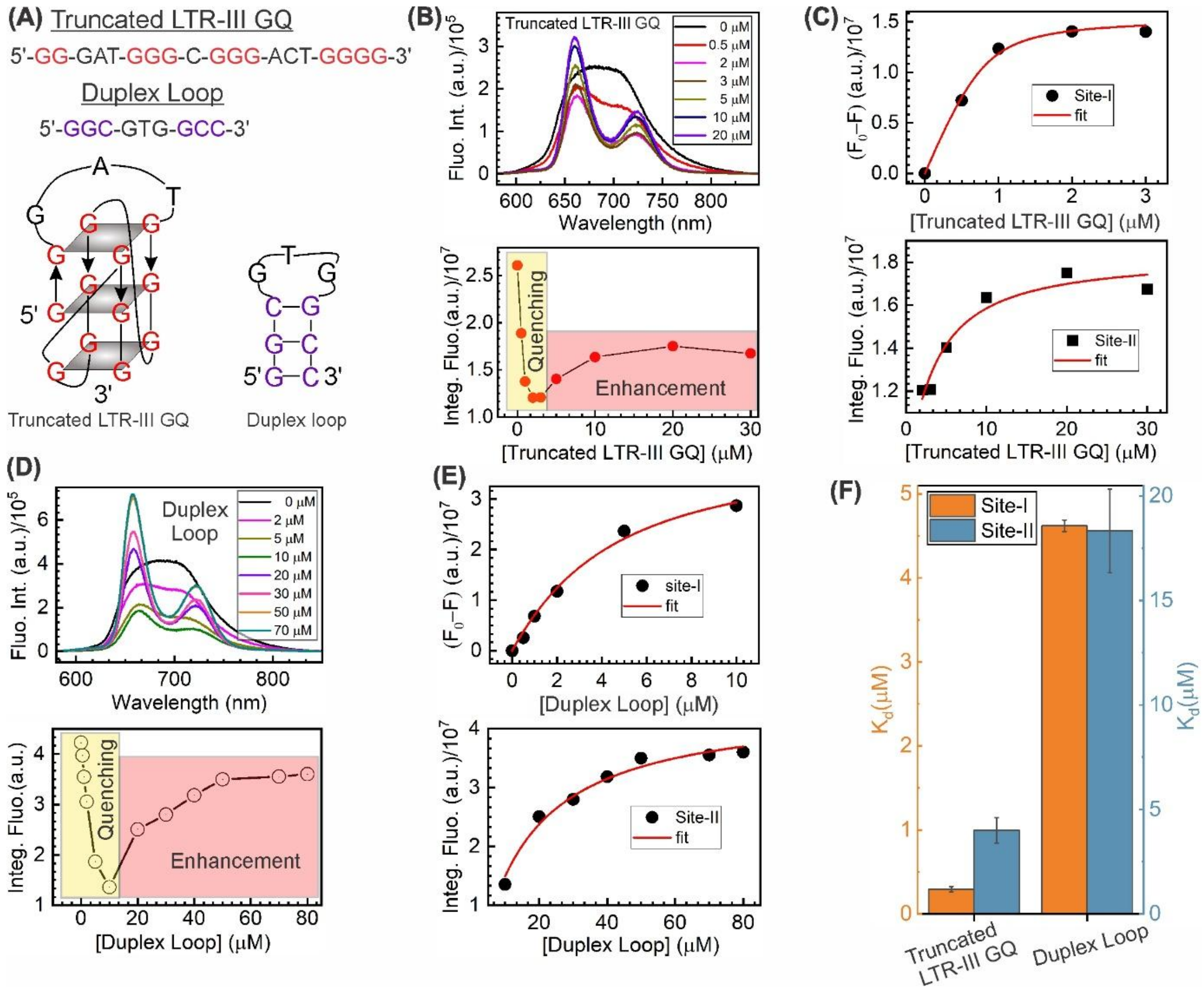


**Figure S2**: **Binding of TMPyP4 to truncated LTR-III G-quadruplex (GQ) and duplex loop constructs.** (A) Sequences and schematic representations of the truncated LTR-III GQ and the isolated duplex loop. (B) Steady-state fluorescence spectra of TMPyP4 (2 μM) upon titration with truncated LTR-III GQ (top) and corresponding integrated fluorescence intensity as a function of GQ concentration (bottom), showing initial quenching followed by fluorescence enhancement. (C) Binding isotherms corresponding to Site-I (quenching regime) and Site-II (enhancement regime) obtained from the truncated LTR-III GQ titration. (D) Steady-state fluorescence spectra of TMPyP4 upon titration with duplex loop (top) and corresponding integrated fluorescence intensity plot (bottom). (E) Binding isotherms for Site-I and Site-II obtained from duplex loop titration. (F) Comparison of dissociation constants ($K_d$) for Site-I and Site-II for truncated LTR-III GQ and duplex loop.

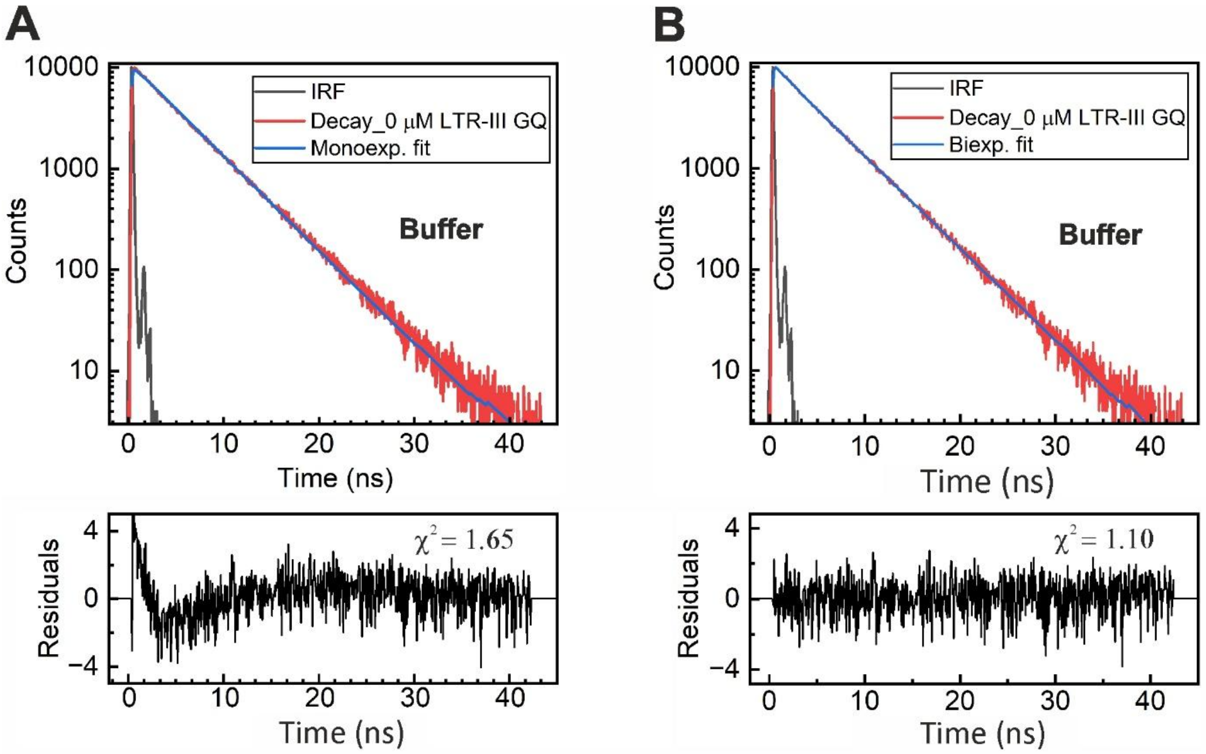


**Figure S3**: Comparison of monoexponential and biexponential fits to the fluorescence decay of TMPyP4 in buffer in the absence of LTR-III G-quadruplex (0 μM GQ). Experimental decay traces are shown in red, fitted curves in blue, and the instrument response function (IRF) in grey. Residuals and reduced chi-square ($\chi^2$) values are shown below each fit.

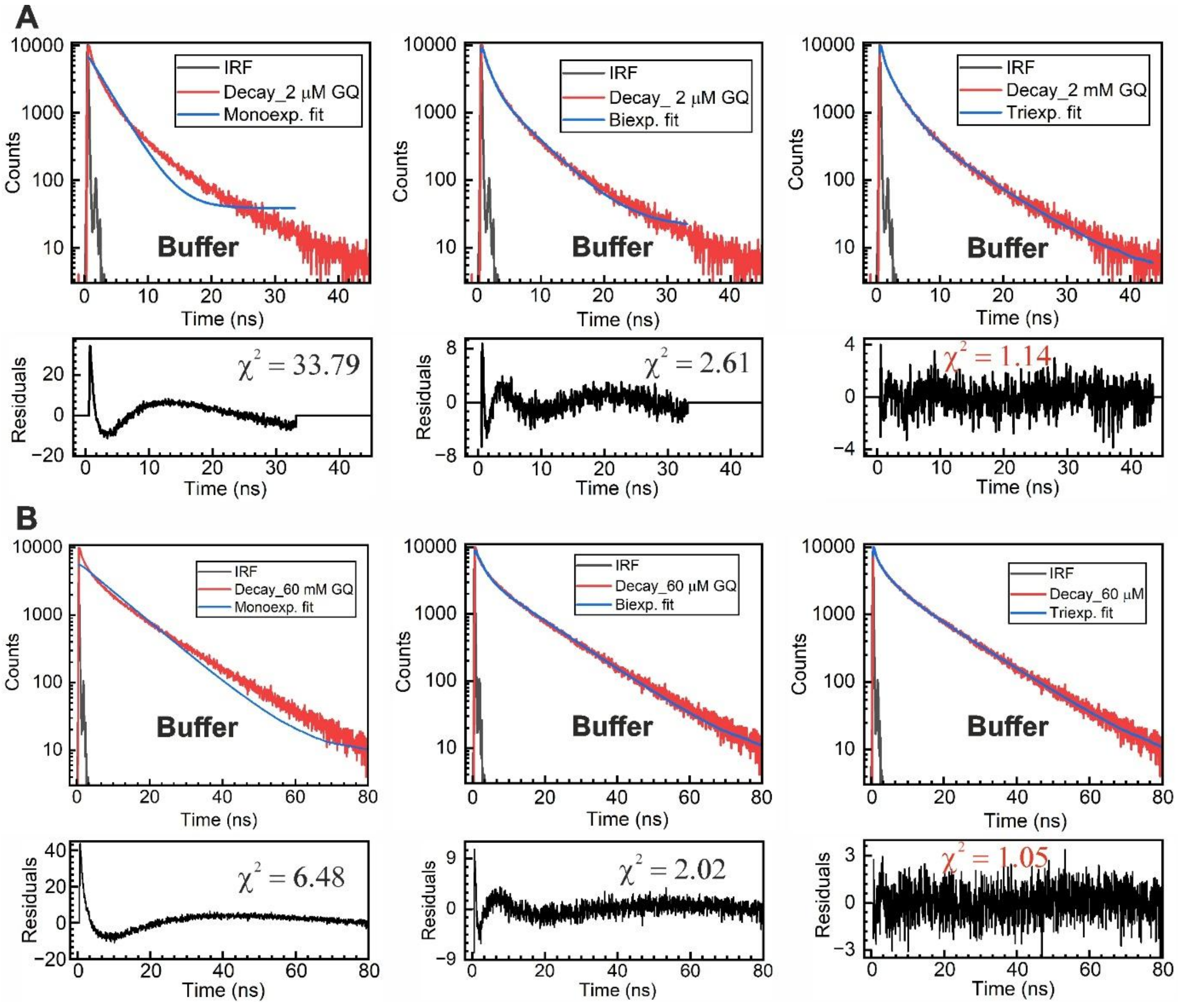


**Figure S4**: Comparison of monoexponential, biexponential, and triexponential fits to the fluorescence decay profiles of TMPyP4 in the presence of LTR-III G-quadruplex (GQ) in buffer at representative concentrations (2 µM and 60 µM). Experimental decay traces are shown in red, fitted curves in blue, and the instrument response function (IRF) in grey. Residuals and reduced chi-square ($\chi^2$) values are shown below each fit. The triexponential model provides the best fit, as evidenced by lower $\chi^2$ values and randomly distributed residuals.

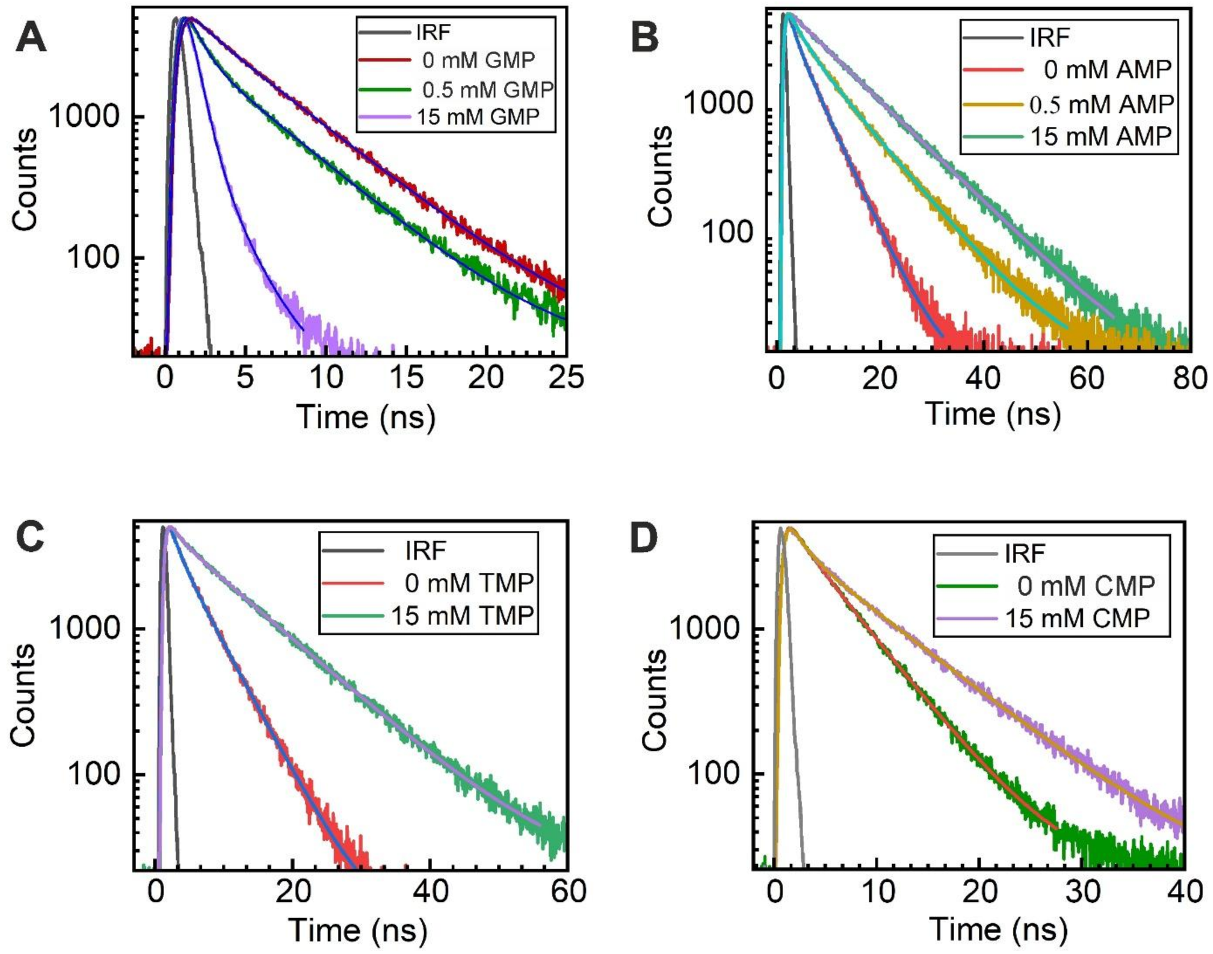


**Figure S5: Time-resolved fluorescence decay profiles of TMPyP4 in the presence of different nucleotides at varying concentrations.** (A) GMP (500 µM and 15 mM), (B) AMP (500 µM and 15 mM), (C) TMP (15 mM), and (D) CMP (15 mM), compared with free TMPyP4. The instrument response function (IRF) is shown in black. Solid lines represent biexponential fits to the experimental data for all traces, except for 15 mM AMP, where a monoexponential fit adequately fits the data. The concentration of TMPyP4 is fixed at 2 µM in all cases.

**Table S1**: TCSPC decay parameters of TMPyP4 (2 µM) in the absence and presence of mononucleotides (GMP, AMP, TMP, and CMP). Except for the 15 mM AMP sample, all fluorescence decays are best fitted with a biexponential model. Here, $\tau_1$ and $\tau_2$ represent the fluorescence lifetime components; $\alpha_1$ and $\alpha_2$ are the corresponding pre-exponential amplitudes; and $f_1$ and $f_2$ denote the intensity fractions associated with each lifetime component.

| **Sample** | **$\tau_1$ (ns)** | **$\alpha_1$** | **$f_1$ (%)** | **$\tau_2$ (ns)** | **$\alpha_2$** | **$f_2$ (%)** | **$\langle\tau\rangle_{amp}$ (ns)** | **$\langle\tau\rangle_{int}$ (ns)** |
|---|---|---|---|---|---|---|---|---|
| TMPyP4 | 1.52 ± 0.14 | 0.11 | 3.79 | 4.79 ± 0.01 | 0.89 | 96.21 | 4.43 ± 0.14 | 4.67 ± 0.01 |
| TMPyP4 + 500 µM GMP | 0.78 ± 0.02 | 0.62 | 22.11 | 4.57 ± 0.03 | 0.38 | 77.89 | 2.21 ± 0.03 | 3.73 ± 0.03 |
| TMPyP4 + 15 mM GMP | 0.58 ± 0.01 | 0.91 | 75.67 | 1.82 ± 0.04 | 0.09 | 24.33 | 0.70 ± 0.04 | 0.88 ± 0.02 |
| TMPyP4 + 500 µM AMP | 4.0 ± 0.01 | 0.50 | 28.60 | 9.98 ± 0.05 | 0.50 | 71.40 | 6.99 ± 0.08 | 8.27 ± 0.03 |
| TMPyP4 + 15 mM AMP | 11.10 ± 0.04 | 1.0 | 100 | | | | 11.10 ± 0.04 | 11.10 ± 0.04 |
| TMPyP4 + 15 mM TMP | 1.31 ± 0.07 | 0.24 | 3.82 | 10.56 ± 0.03 | 0.76 | 96.18 | 8.33 ± 0.07 | 10.20 ± 0.05 |
| TMPyP4 + 15 mM CMP | 1.41 ± 0.05 | 0.38 | 9.97 | 7.90 ± 0.02 | 0.62 | 90.03 | 5.43 ± 0.05 | 7.25 ± 0.03 |

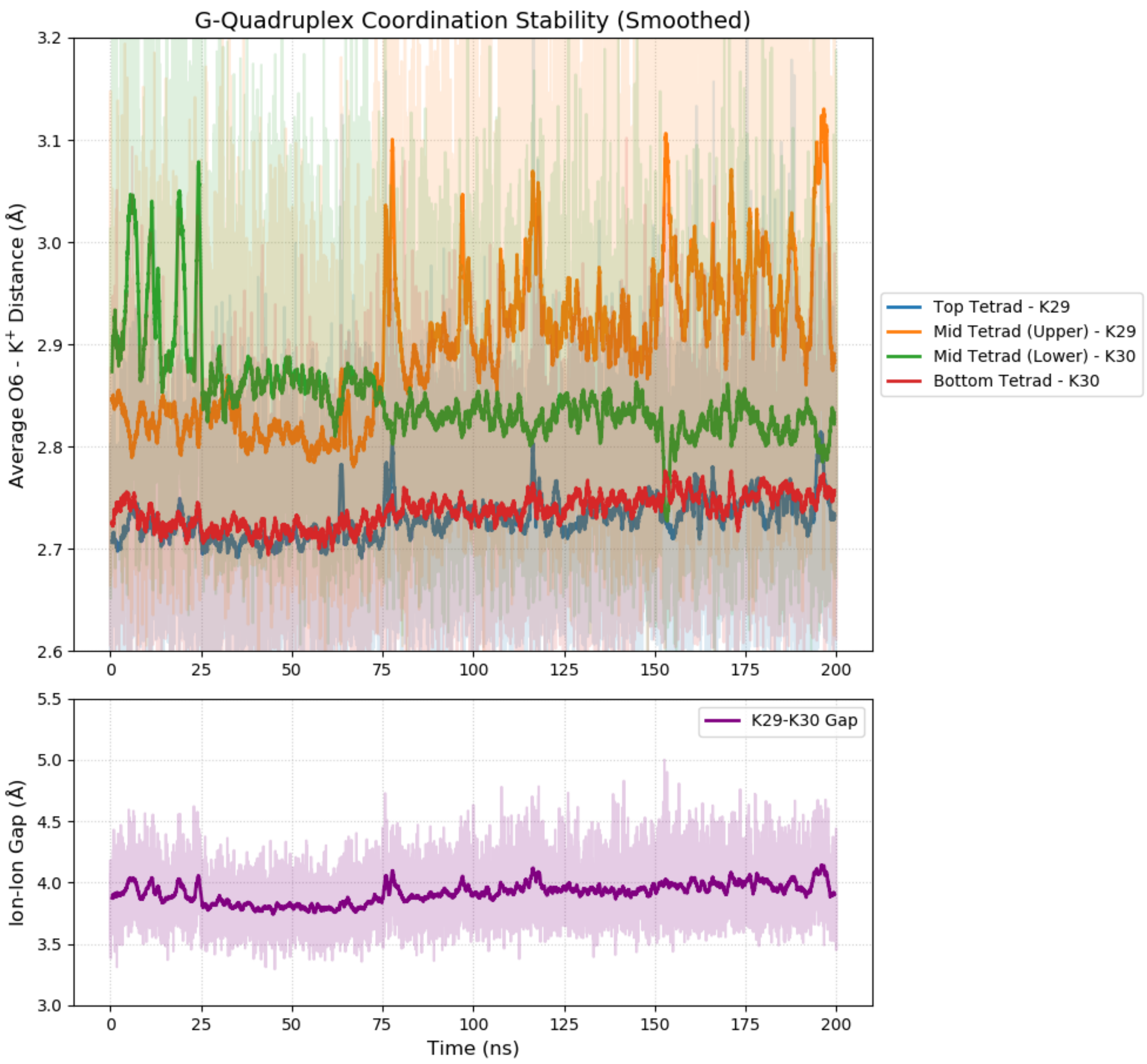


**Figure S6**: **Stability of Channel coordinated K$^+$ ions in the LTR-III GQ. (Top)** Rolling average distance between guanine O6 atoms and the coordinated K$^+$ ions across the three tetrad layers. **(Bottom)** Distance between two channel bound potassium ions, which remains stable at ~3.8 Å throughout the trajectory. Distances were smoothed using a 100-frame rolling average.

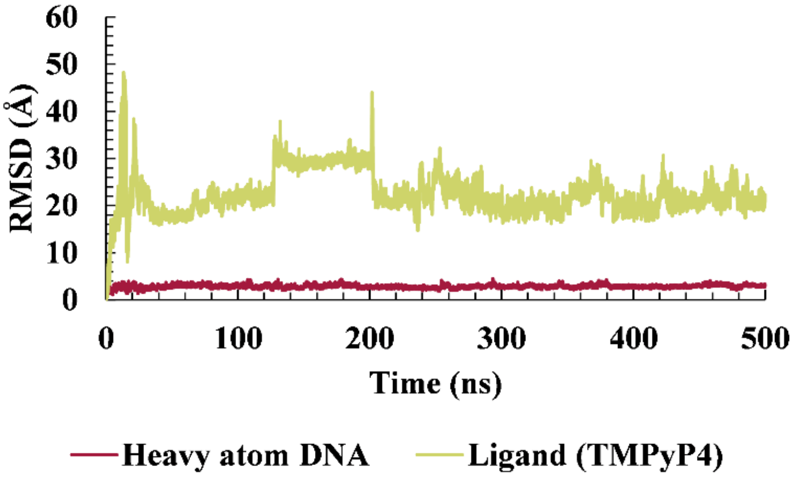


**Figure S7**: **Study of the binding stability of the TMPyP4-LTR-III GQ complex for binding pose at the duplex domain**. RMSD plot of the ligand (TMPyP4) and the DNA heavy atoms over the 500 ns MD simulation for the duplex binding mode.

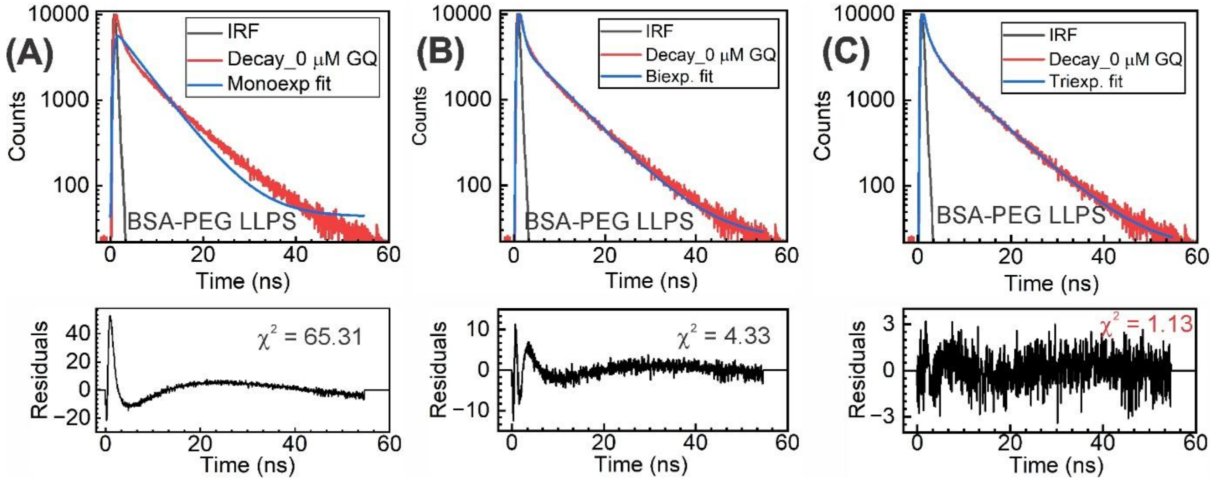


**Figure S8**: **Comparison of (A) monoexponential, (B) biexponential, and (C) triexponential fits to the fluorescence decay profiles of TMPyP4 (2 μM) in BSA-PEG condensates.** Experimental decay traces are shown in red, fitted curves in blue, and the instrument response function (IRF) in grey. Residuals and reduced chi-square ($\chi^2$) values are shown below each fit. The triexponential model provides the best fit, as evidenced by lower $\chi^2$ values and randomly distributed residuals.

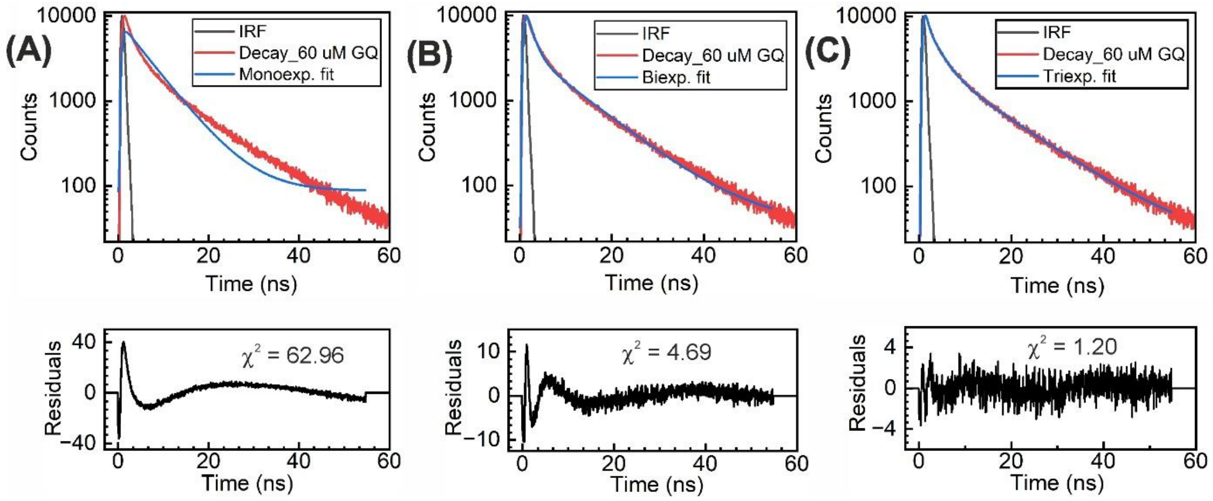


**Figure S9**: **Comparison of (A) monoexponential, (B) biexponential, and (C) triexponential fits to the fluorescence decay profiles of TMPyP4 (2 μM) in the presence of 60 μM LTR-III GQ in BSA-PEG condensates.** Experimental decay traces are shown in red, fitted curves in blue, and the instrument response function (IRF) in grey. Residuals and reduced chi-square ($\chi^2$) values are shown below each fit. The triexponential model provides the best fit, as evidenced by lower $\chi^2$ values and randomly distributed residuals.
